# Was fishing village of Lepenski Vir built by Europe’s first farmers?

**DOI:** 10.1101/2022.06.28.498048

**Authors:** Maxime Brami, Laura Winkelbach, Ilektra Schulz, Mona Schreiber, Jens Blöcher, Yoan Diekmann, Joachim Burger

## Abstract

Today, it is widely accepted that agriculture and settled village life arrived in Europe as a cultural package, carried by people migrating from Anatolia and the Aegean Basin. The putative fisher-forager site of Lepenski Vir in Serbia has long been acknowledged as an exception to this model. Here, the Mesolithic-Neolithic transition - possibly inspired by interaction with the new arrivals - was thought to have taken place autochthonously on site. Our reinterpretation, based on ancient genomes, as well as archaeological and isotopic evidence, revisits this conclusion, indicating that here too, house construction, early village society and agriculture were primarily associated with Europe’s first farmers, thus challenging the long-held interpretation of Lepenski Vir as a Mesolithic community that adopted Neolithic practices. A detailed timeline of population changes at the site suggests that Aegean incomers did not simply integrate into an established Mesolithic society, rather founded new lineages and households. Iron Gates foragers and their admixed descendants appear to have been buried largely separately, on the fringes of the settlement, their diet showing no major shift from aquatic to terrestrial food resources.

## Introduction

One of the most pervasive questions in prehistoric archaeology has long been whether the spread of agriculture into Europe was mediated by migration of new farming groups into the region or adoption by indigenous hunter-gatherer communities. Traditionally cast as a complex and multi-faceted issue, the Neolithic or agricultural transition in Continental Europe is now thought to have primarily involved significant human migration from Anatolia and the Aegean Basin, accompanied in places by low levels of admixture with European Mesolithic populations (Bramanti et al., 2009; Mathieson et al., 2015; Hofmanová et al., 2016). The farmers who migrated to Southeast and Central Europe in the 7th-6th millennia BC brought with them, in addition to plant and animal domesticates, new modes of production and symbolic activities centered on domestic houses and small village communities (Hodder, 1990; Cauvin, 2000; Borić, 2008; Özdoğan, 2011).

While the latest ancient DNA (hereafter aDNA) studies unequivocally show a small Mesolithic contribution to the Danubian Neolithic gene pool, in the order of 2-9% (Hofmanová et al., 2016; Lipson et al., 2017; Mathieson et al., 2018; Nikitin et al., 2019; Marchi et al., 2022), it is not clear how, practically, early farmers and foragers interacted at regional and site levels, nor if these interactions led to Mesolithic communities adopting agriculture and settled village life, as postulated by traditional frontier expansion models in archaeology (Dennell, 1983; Zvelebil & Rowley-Conwy, 1984; Voytek & Tringham, 1990; Zvelebil & Lillie, 2000). An equally plausible scenario in view of the genomic data is that European Mesolithic populations were gradually absorbed into Early Neolithic farming communities without an autochthonous Mesolithic-Neolithic transition (Shennan, 2018). In the Blätterhöhle Cave of Northwestern Germany, for example, foragers and early farmers were found to have lived side by side for 2,000 years without significant shift in subsistence strategies despite clear signals of asymmetric admixture (Bollongino et al., 2013; Lipson et al., 2017).

Long viewed as a missing link between classical Mesolithic and Early Neolithic cultural groups of the Danubian region (Srejović, 1969), Lepenski Vir in today’s Serbia is represented in most archaeology textbooks as an atypical Mesolithic village that successfully exploited its rich and varied ecological niche to provide an alternative path to Neolithic complexity. Stable isotopic analyses conducted since the mid-1990s on skeletal remains from the site indicate that two distinct dietary patterns were present, one based on the exploitation of aquatic resources, the other on a mixture of aquatic and terrestrial resources, possibly reflecting changes in foraging practices and/or the actual introduction of domesticates (**Figure S1**) (Bonsall et al., 1997, 2004). Although changes in subsistence strategies are widely accepted to have come about through interaction with farming communities outside the Gorge, the idea of a local transformation from a predominantly Mesolithic to a predominantly Neolithic culture at Lepenski Vir has remained broadly unchallenged (Borić, 2019, p. 43). As such, the site is ideally suited to revisit processes behind the Neolithic transition in Europe and explore patterns of Mesolithic-Neolithic interactions at a micro-scale, along one of the main corridors of early agricultural dispersal.

Set in the spectacular landscape of the Iron Gates, on the right bank of the Danube, c. 160 km downstream from Belgrade (**Figure 1**), Lepenski Vir is thought to have been reached only by river, or via a narrow path through the mountain (Srejović, 1969, p. 26). It is one of a series of sites that controlled access through the Iron Gates gorge, potentially acting as an important gateway for early farming expansion in Europe. First visited ∼9,500 BC, Lepenski Vir was resettled after a hiatus of at least a thousand years ∼6,200 BC, when the first houses with trapezoidal limestone floors were constructed (**Table 1**) (Bonsall et al., 2002; Borić, 2007, 2016). These semi-subterranean structures (‘pit-houses’), often described as ‘huts’ in the literature, range from 2 to 54 m^2^ in size (mean = 14 m^2^), and display remarkable geometric sophistication, the trapeziform shape being achieved by levelling a platform in the shape of the sector of a circle with the rear end cut into the hillslope (Srejović, 1974, p. 364; Bonsall, 2008, p. 256; Perić & Nikolić, 2011, p. 54). There is still considerable uncertainty regarding the origin and function of the trapezoidal buildings at Lepenski Vir, which recall simpler structures observed in the region, but also display conspicuous Neolithic features, such as intramural burials, elaborate lime plaster floors with thin red or white polished surfaces and a strict internal layout, segmented into zones for cooking and burial activities (Bonsall et al., 2008).

**Figure 1.**
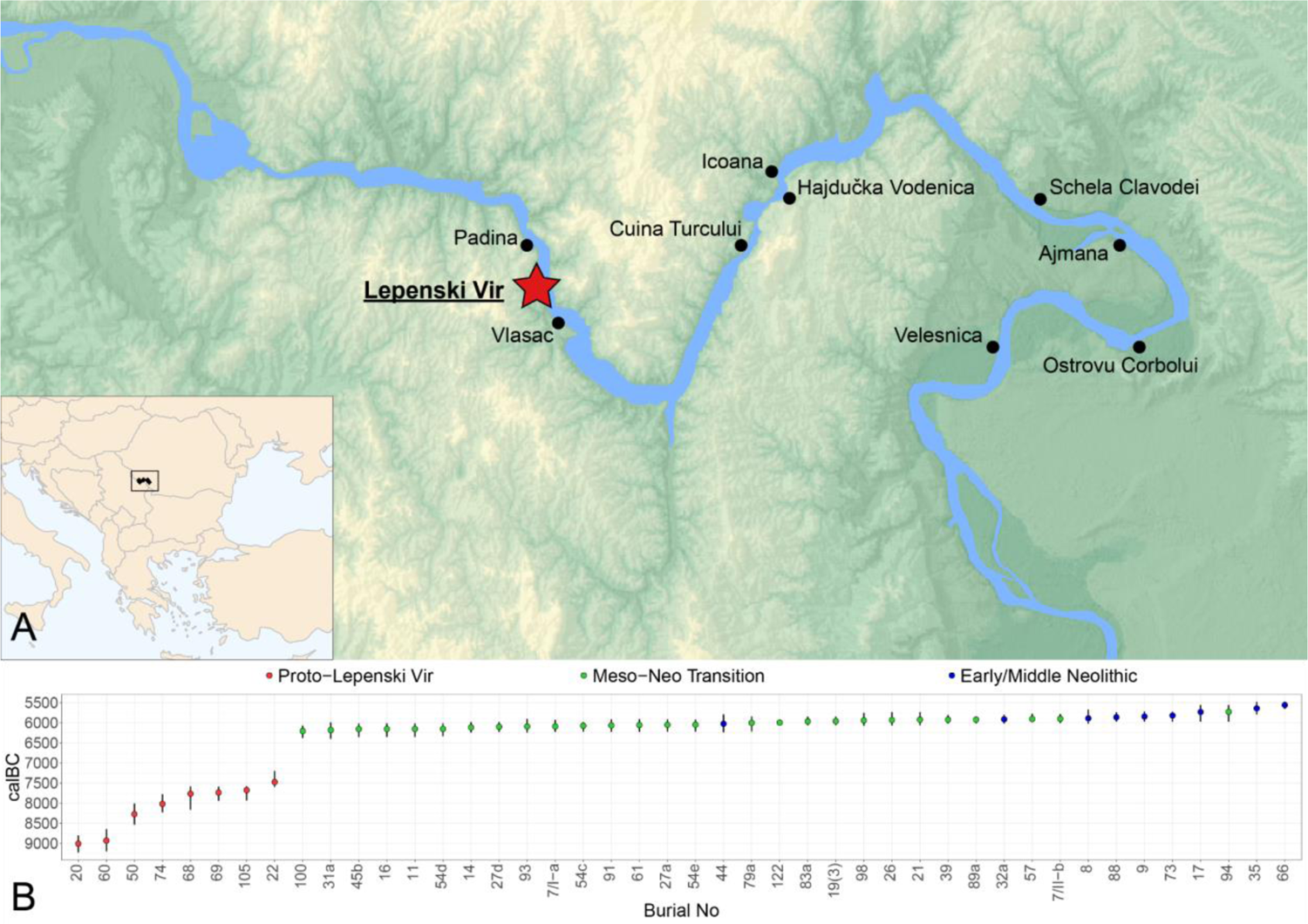
(A) Map of the Iron Gates with the location of key Mesolithic and Neolithic sites including Lepenski Vir; topographic map © EuroGeographics; (B) temporal distribution of ^14^C dated burials of Lepenski Vir (full references in **Electronic Supplementary Material**). Dates corrected for freshwater reservoir effect (Bonsall et al., 2015) and uniformly re-calibrated in OxCal 4.4.2 (Ramsey, 2009) using the IntCal20 calibration curve (Reimer et al., 2020). While Proto-Lepenski Vir and Mesolithic-Neolithic Transition burials are separated by a hiatus, Early/Middle Neolithic ones overlap chronologically with those of the preceding phase.

**Table 1.**
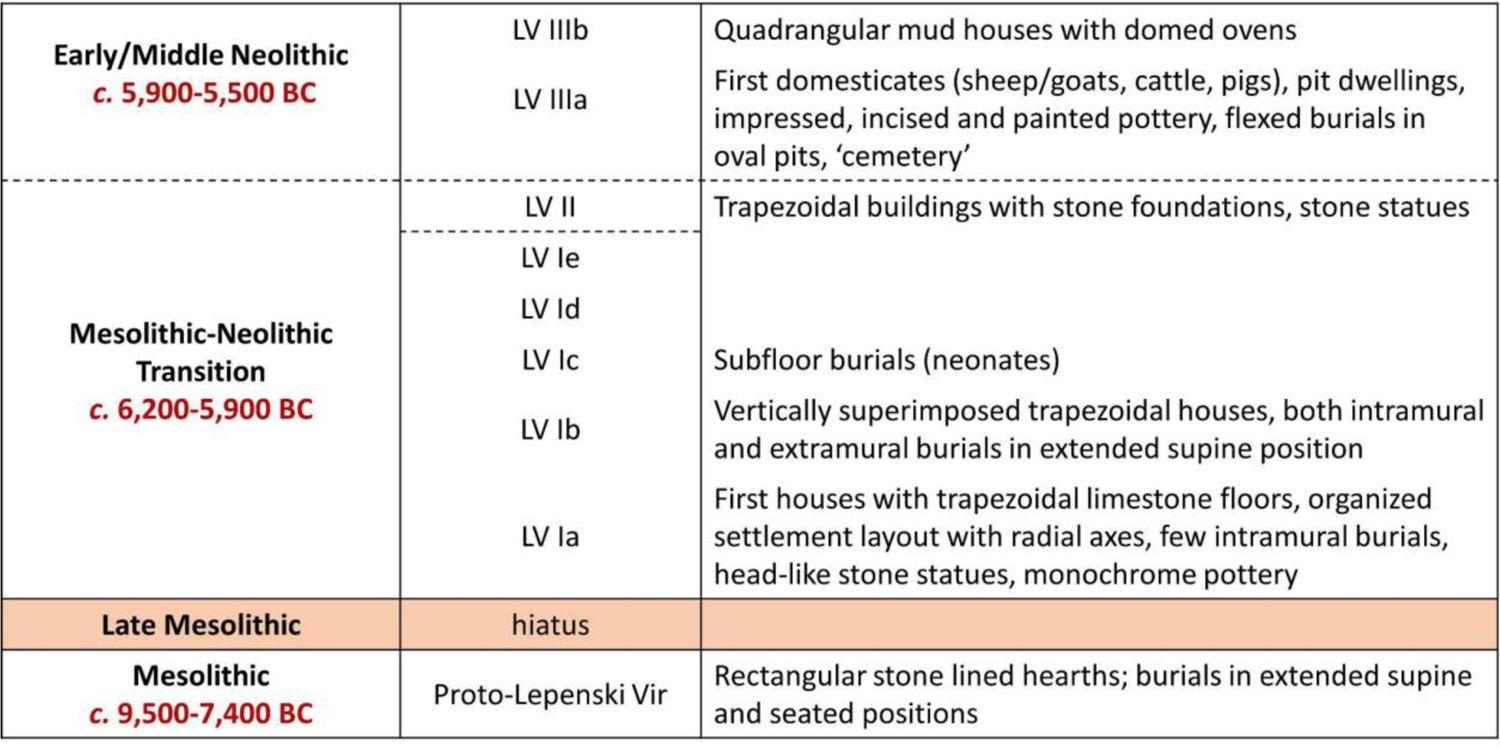
The main phases of occupation at Lepenski Vir and associated features. Phasing after Borić (Borić 2016), with modifications.

|  |  |  |
| --- | --- | --- |
| **Early/Middle Neolithic**<br>**c. 5,900-5,500 BC** | LV IIIb | Quadrangular mud houses with domed ovens |
|  | LV IIIa | First domesticates (sheep/goats, cattle, pigs), pit dwellings, impressed, incised and painted pottery, flexed burials in oval pits, 'cemetery' |
| **Mesolithic-Neolithic Transition**<br>**c. 6,200-5,900 BC** | LV II | Trapezoidal buildings with stone foundations, stone statues |
|  | LV Ie |  |
|  | LV Id |  |
|  | LV Ic | Subfloor burials (neonates) |
|  | LV Ib | Vertically superimposed trapezoidal houses, both intramural and extramural burials in extended supine position |
|  | LV Ia | First houses with trapezoidal limestone floors, organized settlement layout with radial axes, few intramural burials, head-like stone statues, monochrome pottery |
| **Late Mesolithic** | hiatus |  |
| **Mesolithic**<br>**c. 9,500-7,400 BC** | Proto-Lepenski Vir | Rectangular stone lined hearths; burials in extended supine and seated positions |

The abrupt construction of the village as a planned settlement, with rows of trapezoidal houses overlooking the river, coincides with the advent of agriculture in the Central Balkans (**Figure S2**) (Krauß et al., 2018; Porčić et al., 2020; Živaljević et al., 2021). Yet with the exception of the dog, domesticates (sheep, goat, cattle and pig) are not found at the site until the latest phase of occupation, ∼5,900 BC, in the context of disused trapezoidal houses and layers with pit dwellings (Bökönyi, 1970; Borić & Dimitrijević, 2007). Given its location on the steep lower slope of the Koršo mountain, with no adjacent hinterland for cereal agriculture, it is unlikely that Lepenski Vir was ever a food-producing or Neolithic site in a traditional sense. Still, about 20% of the animal bones from Level III belong to domesticated species - older assemblages being too small to categorically rule out animal husbandry (Bonsall et al., 1997, p. 57). Small-scale agriculture, some few kilometres away through the gorge or in the hillsides, has not been proven but remains theoretically possible (Nandris, 1968, p. 67) – no systematic attempts at recovery of plant macro-remains having been made at the now submerged site (Bonsall et al., 1997, p. 57).

For this study, we revisited the interpretation of Lepenski Vir as a permanent settlement of hunters and gatherers in light of recent aDNA evidence for interactions between fisherfolk communities of the Iron Gates and incoming Aegean farmers ∼6,200 BC (González-Fortes et al., 2017; Hofmanová, 2017; Mathieson et al., 2018; Hofmanová et al., 2022; Marchi et al., 2022). Previously reported strontium isotope results seem consistent with a model in which ‘non-local’ Neolithic women, or at least a clear majority of women, intermarried into the native Mesolithic population (**Figure 2**) (Borić & Price, 2013). Such a scenario would result in sex-biased admixture, i.e. predominantly male foragers mating with female farmers, and a predominance of females among Aegean incomers. The forager culture might also be expected to be ‘dominant’ at the site if migrants simply integrated into existing family structures (Borić et al., 2018; Borić, 2019). This is at odds with recent genetic evidence for both male and female incomers at Lepenski Vir (Hofmanová, 2017; de Becdelièvre et al., 2020) and remains to be tested, taking into consideration a broader sample of burials, as well as the age and archaeological context of the burials.

**Figure 2.**
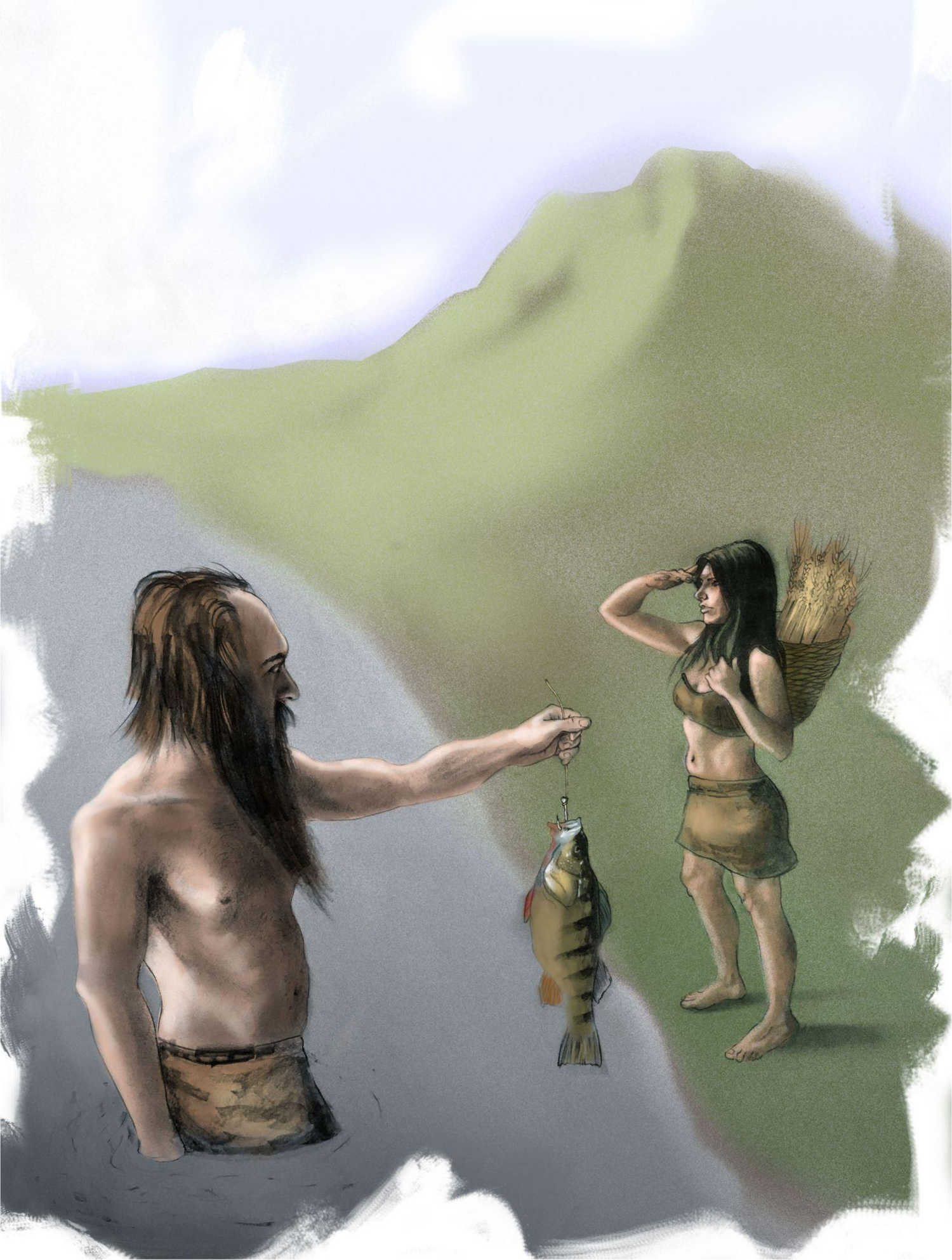
Illustration for the October 2018 issue of Discover ‘When Farmers and Foragers First Met’ (Barna, 2018). Our study shows that women, men and children with an Aegean genetic background settled at Lepenski Vir during the Mesolithic-Neolithic transition phase. Copyright: Prince Parise, reproduced with permission.

To understand who was buried where at Lepenski Vir and how burials relate to the built environment of the site, we re-examined the context of deposition of 34 individuals that have produced autosomal or mitochondrial DNA data (**Table 2**). We compared our results with those obtained at other Iron Gates sites, giving a total of 96 individuals with full, partial and/or mitochondrial genome data to base our analyses on (González-Fortes et al., 2017; Hofmanová, 2017; Mathieson et al., 2018; Hofmanová et al., 2022; Marchi et al., 2022).

**Table 2.** List of previously-reported Lepenski Vir genomes used for this study and summary archaeological, anthropological, chronological and genetic information (for full details, see **Electronic Supplementary Material**). Phasing of burials follows Borić (Borić, 2016), except where new ^14^C dates change interpretation - i.e. Burials 20, 74 (J. Jovanović et al., 2021). Genetic sex reported as XX/XY (where available). Anthropological sex reported as m/f. Key: D: disarticulated; ES: extended supine; FL: flexed. References: (1) Hofmanová 2017; (2) Hofmanová et al. 2022; (3) Mathieson et al. 2018; (4) Marchi et al. 2022.

| Burial No | Genetic ID | Sex | Age | Body position | Cal BC (OxCal $\mu$) | qpAdm | mtDNA | Y | References |
| --- | --- | --- | --- | --- | --- | --- | --- | --- | --- |
| **Mesolithic (c. 9,500-7,400 BC)** |  |  |  |  |  |  |  |  |  |
| 20 | Lepe12 | f | adult | FL | 9006 | .. | U5 | .. | (1), (2) |
| 68 | Lepe51 | XX | adult | ES | 7764 | Iron Gates HG | U4 | - | (1), (2) |
| 74 | Lepe34 | XX | adult | FL | 8017 | .. | U5 | .. | (1), (2) |
| 105 | Lepe47 | XX | adult | D | 7674 | .. | U5 | - | (1), (2) |
| 126 | I5407/<br>Lepe49 | XX | adult | D | .. | Iron Gates HG | H | - | (1), (2), (3) |
| **Mesolithic-Neolithic Transition (c. 6,200-5,900 BC)** |  |  |  |  |  |  |  |  |  |
| 11 | Lepe7 | XX | subadult | ES | 6153 | .. | K | - | (1), (2) |
| 21 | Lepe13 | XY | adult | D | 5921 | .. | X | - | (1), (2) |
| 26 | Lepe15 | m | adult | ES | 5930 | .. | H | .. | (1), (2) |
| 27a | Lepe53 | XX | adult | ES | 6051 | admixed | U5 | - | (1), (2) |
| 27b | Lepe17 | XY | adult | ES | .. | .. | N1a | .. | (1), (2) |
| 27d | Lepe18 | XY | adult | ES | 6099 | admixed | U5 | I2 | (1), (2) |
| 39 | Lepe22 | XX | adult | D | 5921 | .. | J | - | (1), (2) |
| 43 | Lepe23 | .. | subadult | D | .. | .. | K | .. | (1), (2) |
| 54d | Lepe28 | XX | adult | ES | 6151 | .. | K | - | (1), (2) |
| 54e | I4665/<br>Lepe27 | XX | adult | ES | 6047 | Aegean | J | - | (1), (2), (3) |
| 57 | Lepe29 | XX | subadult | D | 5904 | .. | U5 | - | (1), (2) |
| 61 | I4666 | XY | child | ES | 6056 | Aegean | H | R1b | (3) |
| 79b | Lepe37 | XY | adult | D | .. | .. | U5 | .. | (1), (2) |
| 79c | Lepe38 | m? | adult | D | .. | .. | U5 | .. | (1), (2) |
| 82 | Lepe39 | XY | adult | D | .. | Aegean | T2 | C2 | (1), (2) |
| 87(1) | Lepe42 | XY | child | D | .. | .. | U5 | .. | (1), (2) |
| 89a | Lepe44 | XY | adult | ES | 5920 | .. | U5 | .. | (1), (2) |
| 91 | Lepe45 | XY | adult | ES | 6065 | Iron Gates HG | U5 | I2 | (1), (2) |
| 93 | Lepe46 | XX | adult | ES | 6091 | admixed | H | - | (1), (2) |
| 122 | Lepe48 | XY | subadult | D | 5995 | Aegean | K | C1 | (1), (2), (4) |
| **Early/Middle Neolithic (c. 5,900-5,500 BC)** |  |  |  |  |  |  |  |  |  |
| 6 | Lepe3 | .. | subadult | FL | .. | .. | N1a | - | (1), (2) |
| 8 | Lepe6 | XX | adult | FL | 5888 | .. | T1 | - | (1), (2) |
| 17 | I5405 | XX | adult | FL | 5731 | Aegean | HV | - | (3) |
| 19 | Lepe11 | f | adult | FL | .. | .. | K | - | (1), (2) |
| 32a | Lepe20 | XX | adult | FL | 5912 | .. | HV | - | (1), (2) |
| 35 | Lepe1 | XX | adult | D | 5640 | .. | H | - | (1), (2) |
| 66 | Lepe32 | XY | adult | FL | 5559 | .. | N1a | .. | (1), (2) |
| 73 | Lepe52 | XY | adult | FL | 5819 | Aegean | H | G | (1), (2), (4) |
| 86 | Lepe41 | XY | .. | D | .. | .. | J | .. | (1), (2) |

## Results

### Limited gene flow between Aegean and Iron Gates populations in pre-Neolithic times

In line with previous whole-genome studies (Marchi et al., 2022; Mathieson et al., 2018), we find that Iron Gates Mesolithic populations cluster near so-called Western European Hunter-Gatherers (WHG) on a principal component analysis plot, represented for example by Loschbour man in present-day Luxembourg (**Figure 3**). However, a recent study by Marchi et al. has shown that southeastern European hunter-gatherers were separated from the central and western European hunter-gatherers since ∼23 thousand years ago (Marchi et al., 2022; Mathieson et al., 2018). The ancestry of native Mesolithic populations in the Iron Gates is therefore being described throughout this paper as ‘Iron Gates HG’. Aegean and Central Anatolian foragers and early farmers (hereafter ‘Aegeans’), from e.g. Pinarbaşı, Boncuklu and Barcın, represent a distinct ancestry cluster on the PCA (Feldman et al., 2019; Hofmanová et al., 2016; Kılınç et al., 2016; Mathieson et al., 2018). Consequently, in a two-population admixture scenario, individuals from the Mesolithic or Proto-Lepenski Vir phase (∼9,500-7,400 BC) at the eponymous site of Lepenski Vir are well modelled as 100% Iron Gates HGs, without Aegean ancestry that arrives only after 6,200 BC (**Figures 3; S3; S4**). Even though no obvious geographic barrier to gene flow separates the Danube region and the Aegean, which are connected by the Vardar and Morava river systems (Childe, 1929; Nandris, 1970), our analyses confirm that Iron Gates populations were not part of the same mating networks as Aegean and Anatolian populations before the advent of agriculture in the Central Balkans (Mathieson et al., 2018).

**Figure 3.**
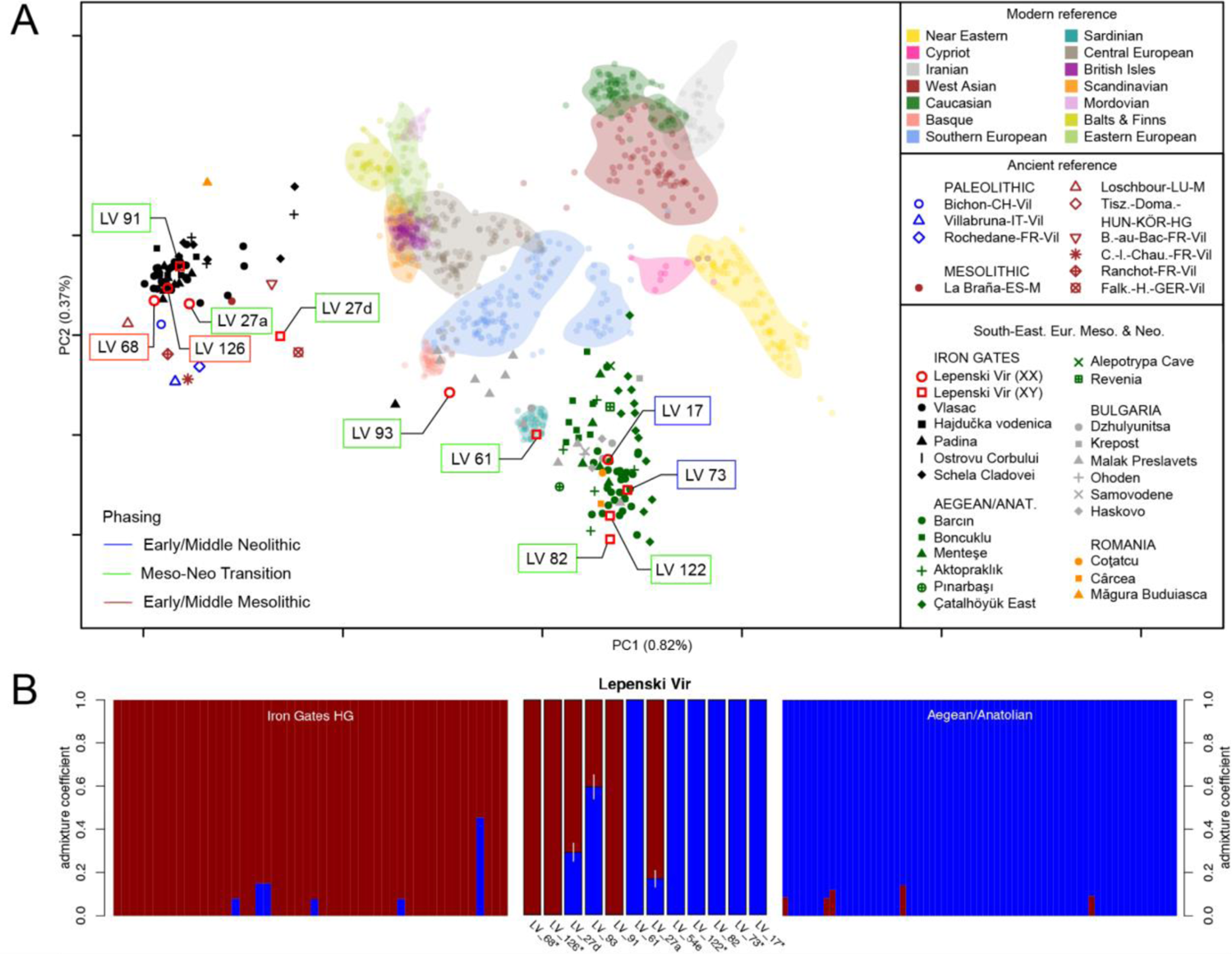
(A) PCA of modern and projected ancient West-Eurasians highlighting the Lepenski Vir (LV) individuals of the different phases; (B) Ancient individuals shown in (A) modelled as mixtures of Iron Gates HG and Aegean-Anatolian ancestry components with qpAdm. Each panel ordered chronologically from oldest to youngest. Lepenski Vir burial No abbreviated as LV.

Several lines of evidence point nonetheless to a small but non-negligible genetic contribution from Aegean to Iron Gates populations in pre-Neolithic times, as reported before (see Sup. Table S2 in (Mathieson et al., 2018)): (1) minimal admixture in five HG individuals from Schela Cladovei and Hajdučka Vodenica (**Figures 3, S3A**), and (2) unusual mitochondrial lineages traditionally associated with Southwest Asian prehistoric populations, namely K1 and H, already present in the Iron Gates respectively since the 10th and 8th millennia BC (**Figure S5**). The significance of these maternal lineages in Mesolithic context depends on the time of admixture between European and Southwest Asian populations, indicating either recent but limited gene flow from the Aegean across the Aegean-Danubian frontier in Mesolithic times, or traces of a more distant admixture process that gave rise to the farming ancestry component, as found by Marchi et al. (Marchi et al., 2022). The latter is the more likely cause for remnants of European Mesolithic-like ancestry inferred in five genomes from Epipalaeolithic Pınarbaşı and Neolithic Boncuklu, Catalhöyük East and Barcın (**Figure S3B**). Regardless, traces of Southwest Asian ancestry in Iron Gates HG populations are too small to account for the abrupt change in ancestry observed ∼6,200 BC at Lepenski Vir, leaving migration as the only plausible explanation.

### Incoming Aegean farmers admixed with local hunter-gatherers but eventually supplanted or fully assimilated them

Patterns of admixture observed among individuals buried at Lepenski Vir (and the neighbouring site of Padina) are consistent with the arrival of communities that are closely related to early farmers of the Aegean Basin during the transitional phase (**Figures 3; S3A; S4B**). Of the six individuals modelled as deriving ∼100% of their ancestry from an Aegean-like source, three are adult or adolescent males (Burials 73, 82, 122) and one is a child of about eight, genetically male (Burial 61) (**Table 2**), suggesting that residence practices and mating patterns were not restricted to Neolithic females moving into Mesolithic communities of the Iron Gates. Even though sample size is small, the fact that residents of all sexes and ages display virtually ‘unadmixed’ Aegean genetic signatures suggests that entire families, as opposed to individual females, relocated to Lepenski Vir after 6,200 BC.

Alternative explanations have been considered to account for the abrupt change in genetic ancestry observed at Lepenski Vir, including the possibility that not all human remains found at the site belonged to people who lived there. A prime contender would be the skull of adolescent male 122 (15-18 years old), which is genetically Aegean-like and was found deposited without a mandible in the packing layer between the floors of trapezoidal houses 47 and 47’, at the centre of the settlement. Much of the iconography at Lepenski Vir, including the fish statues, relates to human heads (Srejović, 1972), suggesting they formed a ritual or funerary focus for the community. Burial 122 may relate to the ritual or funerary deposition of a community member, however, it is also possible that the skull was brought to the site as a trophy or a gift.

At least two of the individuals that have produced genomic information consistent with an Aegean origin (Burials 54e and 61) are primary burials stratigraphically associated with residential floors of trapezoidal houses. Fully articulated skeletons may be expected to belong to people who died at the site or were transported there as corpses shortly after death. Burial 54e lay over stone slabs at the floor level of house 65 and was placed within a stone construction or cairn that was erected upon the trapezoidal house (Bonsall et al., 2008, p. 190; Borić, 2016, p. 510). The burial relates to either the end of the house’s use life or a later re-use. Burial 61, on the other hand, appears to have been inserted through the plaster floor of trapezoidal house 40 (Borić, 2016, p. 512); the fact that the burial was sealed by an anthropomorphic sandstone sculpture at the level of the building floor may suggest that the building was still in use when the burial took place. Burials 54e and 61 were both in extended supine position, confirming their attribution to the Mesolithic-Neolithic transition phase.

We observe both the appearance of new mitochondrial lineages (incl. J, X, T, N1a) and the continuation of existing ones (U5, H) during the Mesolithic-Neolithic Transition phase (**Figure S5**). Two of the individuals that have to be considered as clearly admixed based on the genomic analysis carry U5 maternal lineages (Burials 27a, 27d). Since U5 lineages are rare among Aegean early farmers, it is likely that these individuals had female Mesolithic ancestors, suggesting that local women were part of the mating network of the incoming group that settled at Lepenski Vir ∼6,200 BC. Conversely, the presence of the R1b Y haplogroup in the transitional phase and its low frequency in Aegean individuals (none in 32 [95% CI: 0 - 0.058], **Electronic Supplementary Material**) hints at a HG male contribution to the Lepenski Vir gene pool. Consistent with the latter, we observe that Iron Gates HG males were occasionally buried at the site (Burial 91).

Ancestry proportions estimates for one of the admixed males (Burial 27d) are consistent with 75% Iron Gates HG ancestry on the autosomes and 100% HG ancestry on the X chromosome (**Figure S6**), suggesting that this individual had one Aegean and three Iron Gates HG grandparents. Based on this and the X chromosome of individual 27d being of Iron Gates HG descent, a scenario in which a local HG woman was mating with an admixed male is twice as likely as one in which an admixed female was mating with a HG male. The likelihood of the latter scenario producing the observed data is still 0.5. Assuming the U5 mitochondrial lineage in 27d traces back to a HG woman, admixture must have happened at most two generations in the past.

The picture changes during the latest phase of occupation at Lepenski Vir ∼5,900 BC with the disappearance of the once-prevalent U5 maternal clade and the corresponding rise of pre-existing lineages, such as T, N1a and H, suggesting a disproportional increase of maternal lineages of ultimate Aegean origin (Figure S5). Considering all available ancient mitochondrial genome data from the Iron Gates, haplogroup diversities are lowest in the Mesolithic phase (H=0.38; 95% CI: 0.36 - 0.41), rising considerably in the transitional phase (H=0.73; 95% CI: 0.70 - 0.77), to settle at a high level in Neolithic-period genomes (H=0.81, 95% CI: 0.70 - 0.92) (**Electronic Supplementary Material**).

In a randomly mating population with no survival or fertility differences leading to differential population growth, we expect two base populations with distinguishable ancestry to evolve towards stable ancestry proportions at the individual level over time. Yet, individuals of mixed genomic ancestry are only found in the transitional phase dated to ∼6,200-5,900 BC, whereas later individuals, including adult male 73, are modelled as 100% Aegean. While survival or fertility differences cannot be ruled out, these processes are slow, so that continuous influx of Aegean farmers to the site leading to dilution of Iron Gates HG ancestry over time or even a single replacement event are more likely (**Figure 4**). Interestingly, individual 73 - previously described as ‘local’ (Borić & Price, 2013), albeit with a borderline strontium isotope signature and almost entirely of Aegean ancestry - has a long Run of Homozygosity (∼46cM; **Figure S7**; see also (Marchi et al., 2022)), indicative of recent inbreeding between close relatives (Marchi et al., 2022). Although anecdotal, this might be interpreted as a signal of regulated mating patterns during the last phase of occupation at Lepenski Vir (Level III) despite ‘continuous influx’ of Aegean farmers.

**Figure 4.**
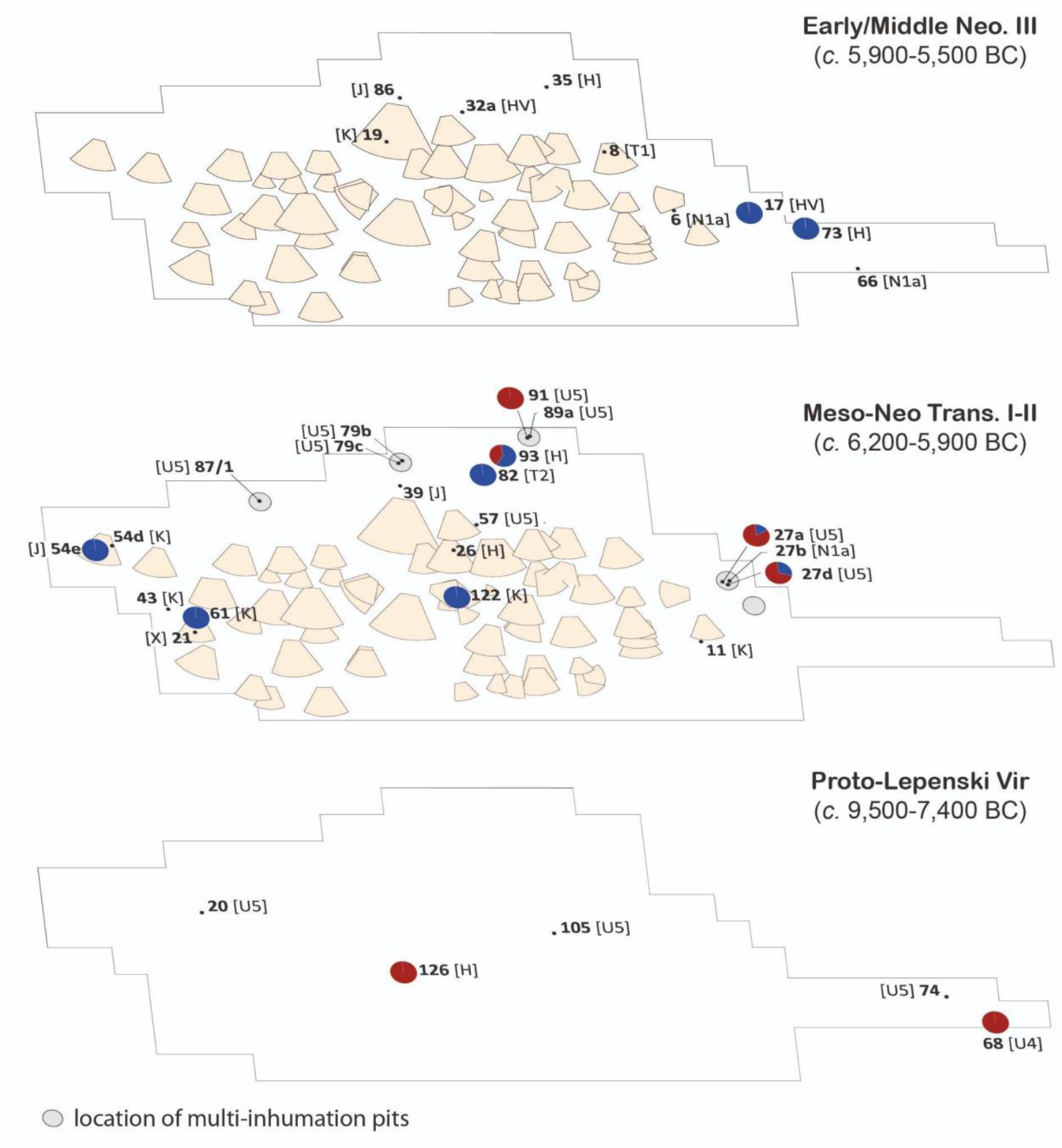
Distribution of ancestry proportions (see Figure 3B; pie charts: red: Iron Gates HG; blue: Aegean-Anatolian) and mitochondrial haplogroups (square brackets) for selected individuals during the three main phases of occupation at Lepenski Vir. While trapezoidal houses were no longer in use during the Early/Middle Neolithic phase, their footprints were still visible and served for burial activities.

### Evidence for structure in the spatial distribution of the burials at Lepenski Vir

We observe structure in the spatial distribution of the burials during the Mesolithic-Neolithic Transition phase, with Aegean-related individuals primarily buried within the habitation space, either inside or in-between houses, while Iron Gates HG individuals appear to be preferentially buried in the upslope area at the rear of the village, in large multi-inhumation pits (**Figure 4**). These pits are unusual features ∼3-4m in diameter, encompassing multiple inhumations, deposited one on top of the other in disarticulated or extended supine position parallel to the Danube, as among Late Mesolithic communities of the Iron Gates (Borić, 2016; Radovanović, 2000). Five such collective burial pits, encompassing Burials 13-16, 27a-f, 89-91, 79(a-c) and 87(1-3), have been identified at Lepenski Vir, one of which (Burial 87) may have had cremations associated with it (Borić, 2016, p. 189).

With mitochondrial haplogroups inferred for as many as ∼20% of known Lepenski Vir individuals, we detect a statistically significant association between U lineages and burials outside in pits (Fisher’s exact test, p=0.00077). To confirm that this association holds at nuclear genomic level, we performed a two-sample permutation test comparing the inferred ancestry proportions between individuals buried inside and outside in pits. The three whole-genomes from the external pits (Burials 27a, 27d and 91) have more Iron Gates HG ancestry on average than other individuals from the transitional phase (permutation test, one-sided p=0.0119).

It is difficult to confirm that areas inside and outside the village were in use at precisely the same time. This is due to the complex stratigraphy of Lepenski Vir, characterized by parallel house sequences that rarely intersect, and the uncertainty of ^14^C dates on human bones, which must be corrected for freshwater reservoir effect in addition to the calibration process (Cook et al., 2001). Summed probability distributions of calibrated radiocarbon dates for burials inside, and outside in pits indicate broad chronological simultaneity, but do not rule out more subtle differences in dating (**Figures S8-S9**). A sequence of overlapping burials with cremations, resembling the multi-inhumation pits encountered at the periphery of Lepenski Vir, has been documented at the nearby site of Vlasac, dating to the 7^th^-6^th^ millennia BC, suggesting continuation of a Late Mesolithic practice in the Neolithic period (Borić et al., 2014).

Based on previously-published stable isotope evidence, it is likely that those individuals buried in pits still consumed high amounts of aquatic proteins, while individuals from the village show a wider range of isotopic signatures, consistent with both a terrestrial diet (plant and/or animal-based) and fishing activities (**Figure 5**). To some extent, body position in the grave reflects the same intra-site variation: Aegean incomers were occasionally buried like Iron Gates HGs in extended supine position, but none of the Iron Gates HGs or U5-carrying individuals identified to date were buried in a flexed position during the transitional phase - a practice observed among Early Neolithic communities of Southeast Europe and among later burials of Lepenski Vir (**Table 2**).

**Figure 5.**
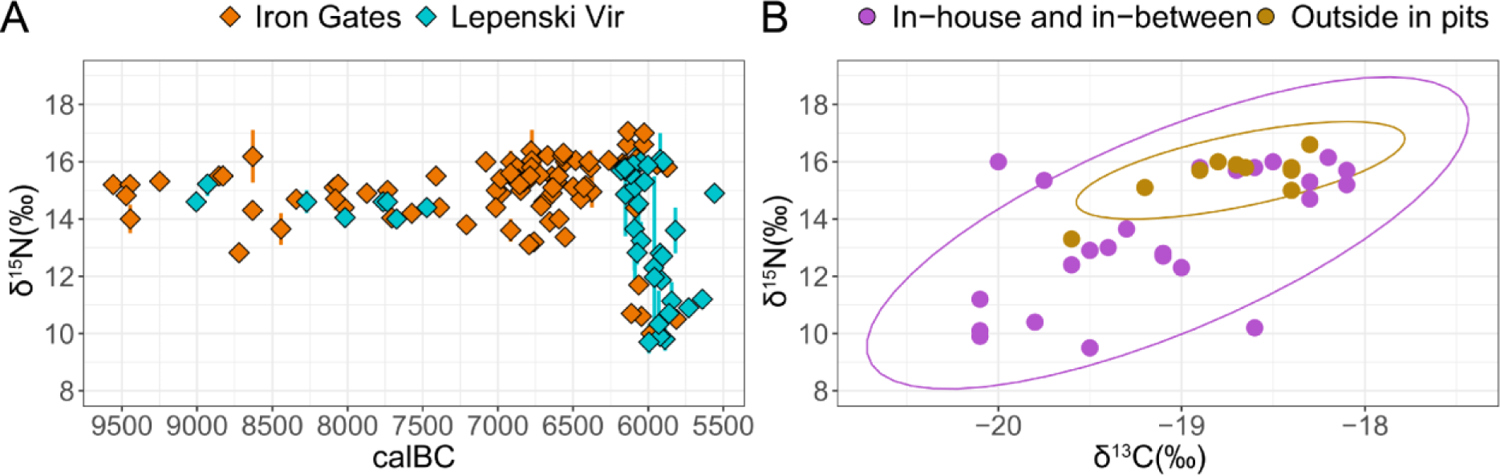
(A) Shift in δ^15^N values observed in directly-dated humans from the Iron Gates (Oxcal μ). Any δ^15^N value above 10‰ is likely to indicate some intake of aquatic proteins; (B) paired median δ^13^C/δ^15^N signatures for individuals of the Mesolithic-Neolithic Transition phase at Lepenski Vir, coloured according to burial location. Ellipses generated with stat-ellipse (assuming a multivariate t-distribution) from the ggplot2 R library. Based on previously reported ^14^C dates and stable isotope values (full references in **Electronic Supplementary Material**).

Taken together, our results suggest that Iron Gates HGs and their descendants were primarily buried on the fringes of the village, maintaining economic and burial practices that were more attuned to a Mesolithic lifestyle. We caution against interpreting our results as deliberate exclusion of Iron Gates HGs from the space of habitation, given limited sample size and the fact that burial location may not reflect where people stayed during their lifetime. Local foragers may have decided to live side by side with incoming farmers, but maintain separate identities with regard to rituals of death and burial. Conversely, Lepenski Vir may have attracted burials beyond the immediate community. Differential access to resources such as domesticated food-plants across multiple generations of inhabitants might highlight conservative dietary traditions, food taboos or even differences in status based on ancestry.

No close genetic relationships between individuals buried at Lepenski Vir could be found that would allow for the reconstruction of familial ties in a biological sense, a fact which may be explained by the lack of systematic sampling of DNA (**Figure S10**). This includes the HGs buried at the periphery of the site in multi-inhumation pits. To explore relatedness beyond immediate kin in a spatial context, we tested for correlation between physical distance among contemporaneous burials at the site and genetic distance measured by outgroup *f_3_* statistics through a Mantel test, but found no correlation (Mantel test, p=0.4143). While these observations remain preliminary, they can be contrasted with those reported for Vlasac, where two neonate brothers (U62 and U69) were buried closely together (Mathieson et al., 2018). Beside this and another previously reported first degree relationship between a father and his daughter (Burials 12 and 24) at Padina, we detect one additional first or second degree relationship between two adult males (Burial 45 and Burial 47) at Mesolithic Vlasac and document a number of additional second to third degree relationships among Iron Gates HGs (**Figure S10**).

It is worth pointing out that non-kinship factors, such as relative age, are likely to have played a role in choice of burial location and burial practice at Lepenski Vir (**Figure S11**), trumping ancestry in the relevant context. Newborn infants (N=41), for instance, were almost without exception buried as foundation deposits beneath trapezoidal house floors during the Mesolithic-Neolithic Transition phase (Stefanović & Borić, 2008). Owing to the liminal status of this age group, it is likely that infants under two months old were afforded different burial rites compared to older children and adults, who were sometimes interred through the floors, i.e. during the use of the buildings or after their abandonment (Radovanović, 2000, p. 340).

Consequently, we expect neonate burials to show different ancestry profiles compared to other community members buried within the houses, who may not have been co-residents. Further DNA analysis shall clarify the position of the neonates within the settlement of Lepenski Vir, and their relationship to adults buried at the site.

## Discussion

### Newcomers were men, women, as well as children born to immigrant parents

Recent biomolecular results are fundamentally changing our image of Lepenski Vir as a Mesolithic village in the process of transition to agriculture. Summing up the evidence at our disposal, we observe that mating was not part of regular interactions between Iron Gates and Aegean populations before the advent of agriculture in the Central Balkans, ∼6,200 BC. Following occasional visits by hunter-gatherers, Lepenski Vir was re-settled by a population consisting of first generation Aegean migrants, Iron Gates HGs and their admixed descendants. By ∼5,900 BC, the incomers’ ancestry became effectively dominant over that of the native Mesolithic population. Unless the site was refounded by another Neolithic community after a gap, this observation is not consistent with the idea of a stable Mesolithic population integrating some Neolithic people and adopting their culture. Only Lepenski Vir and Padina demonstrate significant levels of Aegean ancestry, whereas none of the individuals sequenced to date at Vlasac, including those radiocarbon dated to the 6^th^ millennium BC (Burials H53, U62, U21), show any sign of admixture with Aegean farmers, suggesting that communities stood side by side in the Iron Gates for hundreds of years without regular mating interactions (**Figure S3A**).

Assuming that people buried at the site lived there and were actively involved in its construction, it is likely that Aegean women, men and children settled at Lepenski Vir during the transitional phase (6,200-5,900 BC). A scenario in which Neolithic women married into a Mesolithic group can be rejected, in light of (i) admixture patterns that are not consistent with predominantly male foragers mating with female farmers; (ii) the almost equal representation of females and males among first generation migrants; and (iii) potential evidence for immigrant children (Burial 61). Genomes sequenced to higher coverage are needed to confirm the latter (see **Materials and Methods / *f*-statistics**). At present, our results do not rule out the possibility that Aegean migrants moved to Lepenski Vir as, for example, marriage partners, slaves and/or war captives (Bonsall et al., 2008, p. 194). Should additional genomes indicate, in future, that neonates were born to immigrant parents and their descendants at the site, a case could be made for whole families, as opposed to individuals alone, relocating to Lepenski Vir.

The idea that Lepenski Vir was significantly inhabited during the transition phase by migrant farmers, who traced their ancestry back to the Aegean Basin, is consistent with previously published nitrogen isotope values, which show a shift to lower-trophic level food and presumably cereal-based diets ∼6,200 BC in some humans buried at the site (**Figure S1**). Aspects of the Neolithic Demographic Transition, such as high mortality rates among children below weaning age and population growth, are already visible during the Mesolithic-Neolithic transition phase (Porčić & Nikolić, 2016), suggesting that patterns of replacement observed at genomic level are best explained by continuous gene flow from Neolithic communities outside the Iron Gates and/or higher fertility among individuals with an Aegean genetic background. Interestingly, those individuals with a more terrestrial isotopic signature show the closest genetic proximity to first farmers of Anatolia and the Aegean Basin (Burials 82 and 122). Changes in human isotopic signatures precede the introduction of domesticates at the site, suggesting that newcomers either acquired food from agricultural communities outside the Gorge or came to the site already looking like farmers, isotopically. In all likelihood, they supplemented their diet by fishing in the Danube and raised their children as fishers, hence the wide range of isotopic signatures observed among Aegean farmers (de Becdelièvre et al., 2020).

### Trapezoidal houses at Lepenski Vir associated with incoming Aegean farmers and their descendants

Further questions should be raised about the identification of Lepenski Vir as an established Mesolithic community, in light of evidence for structure in the spatial distribution of the burials during the transitional phase. If Lepenski Vir was indeed a Mesolithic village that integrated a few Neolithic individuals, we might expect members of the community to be buried inside or in close proximity to trapezoidal houses (i.e. their homes). However, individuals assigned to the U clade, who presumably descended from Iron Gates HG populations (or at least had hunter-gatherer or admixed mothers), were all buried, with one exception, some distance away from trapezoidal houses in the upslope area at the rear of the village and show elevated δ**^15^**N isotopic signatures consistent with a fish-based diet (**Figures 5-6**). As things stand, our results speak against a piecemeal adoption of agriculture by Iron Gates HGs and their admixed descendants, who appear to have maintained separate habitation, burial and dietary traditions at Lepenski Vir.

**Figure 6.**
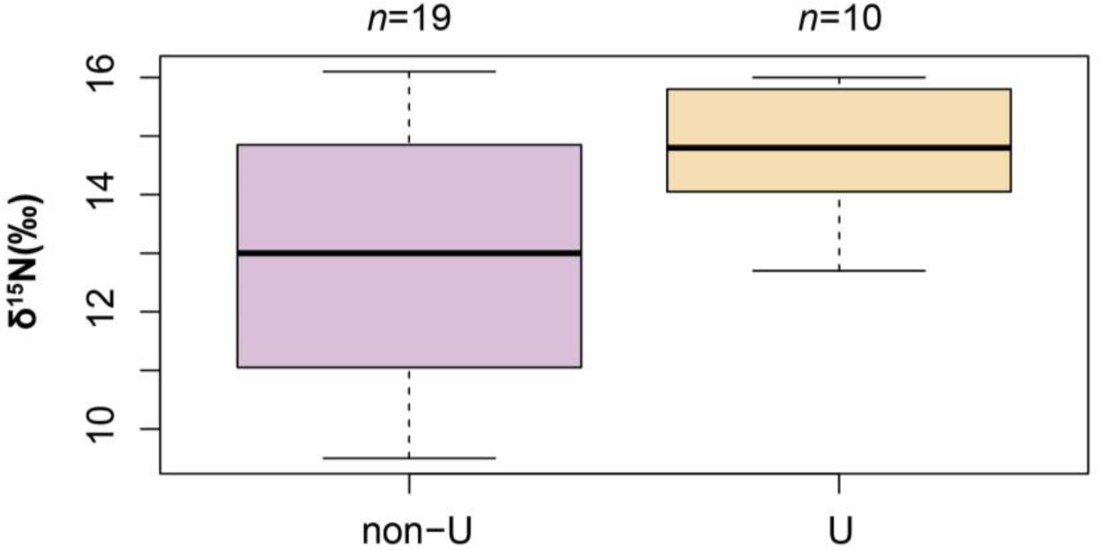
Distribution of median δ**^15^**N values for Lepenski Vir individuals assigned to U and non-U mitochondrial clades (full references in **Electronic Supplementary Material**). The U clade, including U5 and U4, is seen as a marker for European hunter-gatherer ancestry.

While the location was repeatedly visited by foragers, who used it as a burial place and seasonal campsite from at least the 10th millennium BC, the construction of the settlement with the distinctive trapezoidal houses was likely triggered by the arrival of Aegean farmers and their descendants, who were buried inside or in close proximity to houses. Among those individuals that show no signs of admixture with Iron Gates HGs in a two-population qpAdm model, at least three are directly associated with sequences of trapezoidal houses (Burials 54e, 61, 122) (**Figure 3B**). Child 61, who is unlikely to have had more than one hunter-gatherer grandparent if admixed at all (Mathieson et al., 2018), was discovered buried beneath the rear wall of trapezoidal building 40, stratigraphically one of the oldest houses at the site (Bonsall et al., 2002; Borić et al., 2018; Radovanović, 2000).

Given small sample size and lack of systematic sampling strategy within each sequence of trapezoidal houses, we cannot rule out that Aegean incomers were simply reusing a deserted village (Perić & Nikolić, 2016), while the actual inhabitants were those buried outside in pits. Such a scenario appears unlikely in view of the dating of some Aegean-related intramural burials (∼6050 BC), their context of deposition (extended supine position) and the aforementioned stratigraphic relationships with trapezoidal floors and stone statues, but should not be discounted until more genomes are analyzed. Intramural burials are unevenly distributed across the site and rarely exceed three per house, a fact which speaks against a strict interpretation of in-house burials as family graves (**Figure S11**). Trapezoidal buildings may not all have been used as houses. Some may have had specialized functions associated with birth or collective child-rearing, hence the high number of neonate burials associated with them (Stefanović & Borić, 2008).

The association detected in this article between trapezoidal houses and Aegean ancestry at Lepenski Vir adds a new twist to the long-standing debate regarding the origins of this house form in the Iron Gates (Bonsall et al., 2008). Initially described as an autochthonous Mesolithic phenomenon (Srejović, 1972), trapezoidal houses were subsequently regarded as a marker of hybridization between Mesolithic and incoming Early Neolithic cultural groups, pottery being found within some of the trapeziform-shaped houses at Padina (B. Jovanović, 1969, 1972). Starčevo-like Early Neolithic pottery, imported Balkan flint and ground edge tools were later reported from trapezoidal house floors of Lepenski Vir, including a complete pot in the ‘ashplace’ of House 54 (Bonsall et al., 2008, p. 192; Garašanin & Radovanović, 2001).

Further research on the Mesolithic of the Iron Gates has indicated that small trapeziform-shaped pit structures, occasionally associated with rectangular stone-bordered hearths, were already being constructed at Vlasac, Schela Cladovei and other Late Mesolithic Iron Gates sites (Bonsall, 2008, p. 256; Borić et al., 2008, p. 278, 2014; Boroneanț et al., 2014; Rusu, 2011, 2019). These structures fit within a local tradition of increased sedentism and open-air settlements on the Danube starting ∼7,600 BC (Bonsall, 2008, p. 240), their unusual shape being thought to reference the trapezoidal Treskavac mountain on the other side of the river at Lepenski Vir (**Figure S12**). As larger trapezoidal structures with a broader end facing the river appear to be predominantly associated with layers that are Early Neolithic in date (Bonsall, 2008, p. 256), a fusion between a local building form and the habitation practices of incoming farmers seems to have taken place - Lepenski Vir being the most spectacular product of that syncretic process. Trapeziform-shaped pit structures with their back against the hillslope may have been ideally suited for the gorge environment, which was prone to mudslides, as indicated by hillwash deposits at Vlasac and other Iron Gates sites (Borić et al., 2008).

The arrival of Aegean farmers in the Iron Gates likely led to renewed elaboration and construction of systematically-planned villages such as Lepenski Vir, which displays typical Early Neolithic practices, including subfloor burials and vertical superimposition of houses, leading to *tell*-like accumulation of deposits by the side of the mountain (Borić, 2008; Chapman, 2000). Burials occurred within a symbolic environment that was not unlike that observed at early agricultural sites, with distinct areas reserved for burial activities at the rear of the houses, away from the central fire installation (Hodder, 1990). The pyrotechnology needed for making elaborate red limestone floors has good parallels in Neolithic Anatolia and reflects what John Nandris has described as a “Neolithic mode of behaviour” (Nandris, 1988, p. 14).

If Iron Gates HGs maintained separate traditions, as suggested here, how to explain the ubiquity of Mesolithic practices at Lepenski Vir? The hybrid character of the site is generally well accepted, with the forager culture seen as “dominant” during the transitional phase (Borić, 2019, p. 43). Assuming that burial in extended supine position parallel to the Danube with the head pointing downstream was the norm among Late Mesolithic Iron Gates HGs (Bonsall & Boroneanț, 2018), then some of the incoming Aegeans can be considered to be buried in a similar fashion to local people during the transitional phase (e.g. Burials 61, 54e). Conversely, not a single Iron Gates HG was found buried in a typically Aegean Neolithic manner - the only U5-carrying individuals displaying a flexed position (Burials 20, 74) having been recently re-dated to the 10th and 9th millennia BC (J. Jovanović et al., 2021). What we are seeing is a brief ‘window’ in the development of farming, where farmers become foragers when confronted with well-established local traditions and subsistence practices. Importantly, this suggests that agriculture is not quite as robust a tipping point as usually assumed. Those individuals identified genetically as ‘Aegeans’ display a remarkable ability to adopt and transform local practices, confounding expectations of a unidirectional shift from Mesolithic to Neolithic societies.

## Conclusion

A model of Mesolithic-Neolithic transition for inland Europe hinging on Lepenski Vir as a Mesolithic village that transitioned to agriculture is not sustainable. Even in a remote environment like the Iron Gates, the influence of Europe’s first farmers was pervasive. The arguments outlined in this article do not in any way challenge the exceptional nature of Lepenski Vir, which remains the only Early Neolithic-period site in Europe with evidence of recent admixture (two generations or less) between early farmers and foragers. Especially noteworthy is the readiness of Aegean farmers to adopt local Mesolithic burial and dietary practices, a fact which may be contrasted with the more conservative outlook of sampled Iron Gates HG individuals. Lepenski Vir clearly differs from traditional Early Neolithic communities of Southeast and Central Europe, which show little if any influence from the Mesolithic world. Why Aegean farmers chose to settle in a location that was not well suited for cereal agriculture is an intriguing question, which will no doubt come under sharp scrutiny in light of our re-interpretation of the site. At Lepenski Vir, the wild and the domestic were never very far.

## Materials and Methods

### Bioinformatics pipeline

#### Alignment

Residual adapter sequences were trimmed from the FASTQ files with trimmomatic (Bolger et al., 2014) with corresponding Illumina adapter sequences, filtering for a minimal length of 30bp after trimming. Forward and reverse read-pairs were subsequently merged with BBDuk (Bushnell et al., 2017), only retaining reads with a minimal overlap of three base-pairs. The trimmed and merged FASTQ files were aligned to the human reference genome (GRCh37/hg19) using bwa aln (Li & Durbin, 2009). During the conversion to BAM format with samtools (Li et al., 2009), only reads with a mapping quality >= 30 were kept. PCR duplicates were removed with sambamba (Tarasov et al., 2015) with default parameters. Prior to the variant calling, the alignments were re-aligned around known SNPs and InDels using GATK version 3.7 (McKenna et al., 2010). DNA authenticity was assessed based on the mitochondrial chromosome using ContamMix (Fu et al., 2013).

#### Variant Call

All variant calling was carried out following the approach described in Hofmanová et al. (Hofmanová et al., 2016), by using the ATLAS package ((Link et al., 2017) commit=647daf7). Majority-allele calls (method=majorityBase) were produced for the SNPs overlapping the 1240K capture array (Mathieson et al., 2015), as well as for the mitochondrial and the Y-chromosome. Diploid Genotype calls were obtained with the maximum likelihood approach described in Hofmanová et al. 2016 (method=MLE) for sites found to be bi-allelic in the 1000 Genomes data-set (1000 Genomes Project Consortium et al., 2010).

#### Uniparental markers and molecular sexing

The molecular sex of each sample was determined following the approach described in Skoglund et al. (Skoglund et al., 2013), Y-chromosomal haplogroups were assigned from BAM files using Y-leaf (Ralf et al., 2018). For MT-haplogroup determination majority-allele calls from each MT-chromosome were uploaded to haplogrep2 (Weissensteiner et al., 2016). Haplogroup diversity *H* was estimated as:

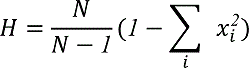

based on the haplotype frequency *x*_i_ in *N* samples. 95% confidence intervals were obtained via leave-one-out jackknife resampling.

### Population genetics

#### Principal Component Analysis

PCA was performed with LASER version 2.04 (Wang et al., 2014) following the approach described in Hofmanová et al. (Hofmanová et al., 2016). After generating a reference space of modern Eurasian individuals (data published as part of reference (Lazaridis et al., 2016); see Supplementary Methods for list of populations), we projected BAM files (see Supplementary Methods for list of individuals) onto the reference space via Procrustes analysis implemented in LASER averaged over ten replicates, using pileup files generated with Samtools version 1.9 (Li et al., 2009) with the suggested filter criteria of minimum mapping quality 30 and minimum base quality 20.

#### *f*-statistics

All *f*-statistics, i.e. outgroup *f*_3_ and *f*_4_ admixture proportions, were computed with *qp3Pop* and *qpAdm* from the ADMIXTOOLS (Patterson et al., 2012) package respectively, with default parameters and on the sites defined by the *HOIll* set of SNPs (Lazaridis et al., 2016). All *qp3Pop* analyses are performed with Khomani as outgroup, all *qpAdm* runs used the set of outgroups Han, Karitiana, Mbuti, Onge, Papuan, Mota, Ust’-Ishim, MA1, El Miron, GoyetQ116-1. Pseudohaploid genomes were retrieved from David Reich’s lab website (https://reich.hms.harvard.edu/allen-ancient-dna-resource-aadr-downloadable-genotypes-present-day-and-ancient-dna-data; downloaded January 20, 2021); in addition, we used our own majority allele calls for the individuals from Yaka et al. (Yaka et al., 2021) as described in section **Bioinformatics pipeline / Variant Call**. See **Supplementary Methods** for the list of individuals used to represent the Iron Gates hunter gatherer and Aegean/Anatolian farmer ancestry components.

Note that due to different sets of source individuals our ancestry proportions may differ from previously published ones. For example, individual LV_61 (I4666) has originally been described as a mixture of north-western-Anatolian-Neolithic-related and hunter-gatherer-related ancestry (Mathieson et al., 2018). We find no statistical support for a two-component model over a simpler exclusively Neolithic one, although both fit well (mixture of 9% Iron Gates hunter gatherer and 91% Aegean/Anatolian farmer: tail probability 0.85; pure Aegean/Anatolian farmer: tail probability 0.88, *p*-value for nested model 0.52). This may be explained by the relatively small number of 44,695 SNPs available for the analysis of LV_61.

#### X/Autosomal ancestry

For the comparison of ancestry patterns on X and autosomes, individual ancestries were estimated based on the *HOIll* set of SNPs (Lazaridis et al., 2016). Following (Oteo-García & Oteo, 2021), we quantified hunter-gatherer ancestry by computing *f_3_*(Aegean, Test, HG) / *f_2_*(Aegean, HG) separately for the X chromosome and autosomes. Here, ‘Aegean’ and ‘HG’ stand for the same sets of samples used in the *qpAdm* analysis to represent Aegean/Anatolian farmer and Iron Gates hunter gatherer ancestry, respectively. 95% confidence intervals were obtained by re-sampling, leaving out SNPs in 5 centimorgan windows.

#### Kinship analyses

Biological relatedness of the Iron Gates samples was determined following the methodology outlined in Yaka et al. (Yaka et al., 2021), using READ (Monroy Kuhn et al., 2018), lcMLkin (Lipatov et al., 2015) and ngsRelate (Hanghøj et al., 2019; Korneliussen & Moltke, 2015). For the READ analysis majority-allele calls overlapping the 1240K capture array (Mathieson et al., 2015) were used. For lcMLkin and ngsRelate, phred-scaled genotype likelihoods from the MLE-calls were used as input. All analyses were repeated three times on three different datasets. First for all samples jointly, second for all samples with predominantly hunter-gatherer ancestry, as well as admixed individuals, third with all samples with predominantly farmer ancestry, as well as admixed individuals. Admixture proportions were taken from the *qpAdm* analysis described above. The merged MLE-calls were pruned, based on linkage disequilibrium with plink (--indep-pairwise 200 25 0.4, (Purcell et al., 2007)) and only polymorphic SNPs with a frequency >= 5% were kept, which were present in at least 30% of the samples. For ngsRelate and lcMlkin, only results for sample pairs with at least 5,000 overlapping SNPs were considered.

### Radiocarbon dating

For an overview of the radiocarbon dates considered for this project see **Electronic Supplementary Material**. All dates were uniformly re-calibrated in OxCal 4.4.2 (Ramsey, 2009) using the IntCal20 calibration curve (Reimer et al., 2020). Correction for freshwater reservoir effect follows Bonsall et al. (Bonsall et al., 2015): for human bones with δ*^15^N* ≥ *8*.*3‰*, uncalibrated dates of the form μ ± σ were adjusted to 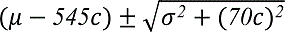 where 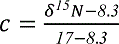.

Summed probability distributions of calibrated radiocarbon dates were calculated in OxCal using the Sum() function.

### Statistical analyses

Fisher’s exact test was run in R with the command ‘fisher.test’, and the permutation test to compare ancestry proportions inside and outside in pits was performed with ‘permTS’ from the ‘perm’ R library. The Mantel test for correlation between outgroup *f_3_* and spatial distance was performed with the command ‘mantel.rtest’ from the ade4 R library. Note that a Mantel-test assumes a smooth correlation, which may not be the case on the *f_3_* scale, especially given the admixture scenario. The two-sample Kolmogorov-Smirnov test comparing the summed probability distributions of calibrated radiocarbon dates (*n_1_* = *8*, *n_2_* = *7*) at significance level α = *0.05* was implemented in R by testing if the maximal absolute difference between the cumulative probability distributions exceeds 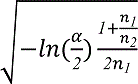. The credible interval for the R1b Y-haplogroup frequency given no R1b in *n* observed male individuals was computed as the 95th quantile of *Beta(0.5, 0.5 + n*).

## Supporting information

Electronic Supplementary Material

## Acknowledgements and funding sources

M.B. was supported by an Individual Fellowship from the H2020 Marie Skłodowska-Curie Actions (793893) and by the Institut National de Recherche Archéologique, Luxembourg. Y.D. is supported by the ERC Advanced Grant 788616 “Yamnaya Impact on Prehistoric Europe” (YMPACT) awarded to Volker Heyd.

Parts of this research were conducted using the supercomputer Mogon and/or advisory services offered by Johannes Gutenberg University Mainz (hpc.uni-mainz.de), which is a member of the AHRP (Alliance for High Performance Computing in Rhineland Palatinate, www.ahrp.info) and the Gauss Alliance e.V.

We thank Bogdana Milić for helpful information and Katie Meheux, David Shankland and Stephen Shennan for comments on a preliminary draft. Prince Parise kindly granted permission to reproduce the Discover illustration.

## Supplementary Methods

### Principal Component Analysis

Groups consist of the following populations: Southern European (Italian North/South, Sicilian, Spanish/-North, Canary Islander, Maltese, Greek), Basque, Sardinian, Cypriot, Central European (Albanian, Bulgarian, Romanian, Hungarian, Croatian, Czech, German, French), Eastern European (Russian, Ukrainian, Belarusian, Polish, Sorb), Mordovian, Baltic and Finnish (Estonian, Lithuanian, Finnish), British Isles (English, Orcadian, Scottish, Irish/- Ulster, Shetlander), Scandinavian (Icelandic, Norwegian), Caucasian (Georgian, North Ossetian, Abkhasian, Chechen, Adygei, Lezgin, Kumyk, Balkar), West Asian (Turkish, Armenian), Iranian/-Bandari, Near Eastern (Palestinian, Druze, Jordanian).

We projected BAM files for ancient reference individuals Bichon-CH-UP (E. R. Jones et al., 2015) (Upper Paleolithic [UP] from Switzerland [CH]), Villabruna-IT-Vil (Fu et al., 2016) (UP, part of the the Villabruna [Vil] cluster, from Italy [IT]), Rochedane-FR-Vil REF (UP, Vil cluster, from France [FR]), La Braña-ES-M (Olalde et al., 2014) (Mesolithic [M] from Spain [ES]), Loschbour-LU-M (Lazaridis et al., 2014) (from Luxembourg [LU]), Tisz.-Doma.-HUN-KÖR-HG (Gamba et al., 2014) (Mesolithic hunter-gatherer [HG], Körös [KÖR] archaeological context, from Tiszaszölös-Domaháza, Hungary [HUN]), B.-au-Bac-FR-Vil REF (Le Vieux Tordoir at Berry-au-Bac), C.-l.-Chau.-FR-Vil REF (Les Fontinettes at Cuiry-lès-Chaudardes), Ranchot-FR-Vil REF, Falk.-H.-GER-Vil REF (Falkensteiner Höhle, Germany [GER]).

The list and references of South-Eastern European Mesolithic and Neolithic individuals additionally projected can be found in the **Electronic Supplementary Material**.

#### *f*-statistics

The admixture components used for *qpAdm* are Iron Gates hunter gatherers (LEPE51, VLASA4, VLASA10, VLASA37, VLASA41, VLASA44, I4660, I4871, I4872, I4873, I4875, I4876, I4877, I4878, I4880, I4915, I4916, I5233, I5234, I5235, I5236, I5237, I5238, I5239, I5242, I5244, I5401, I5407, I5771, I5772, I5409_published, I4881_published, I4914_published, I5773_published), and Aegean/Anatolian farmers (I3879, I0698_published, I0704_published, I3498, I5427, I2937, I1508, AKT6g, Bar8, CCH285, CCH294, CCH311, cth006, Lepe40, I0676, I2533, I5405, Bon001.SG, Bon002.SG, I0707, I0726, I0736, I0744, I0745, I0746, I1096, I1097, I1099, I1579_published, ZHAG_BON004.A0101_Luk10, ZHJ_BON024.A0101_Luk84).

See Electronic **Supplementary Material** for references.

#### MT-tree

To build a tree from the mitochondrial DNA sequences, BAM files for all available samples were downloaded. Calls for the mitochondrial genomes were done with ATLAS (Link et al., 2017) using the majorityBase option. During calling the first 3 bases of each read were ignored to minimize the potential impact of post-mortem deamination. Consensus sequences for all individuals were extracted from the VCFs and pairwise aligned using MAFFT (Katoh et al., 2002) with default options. The maximum likelihood tree was built from the pairwise alignment with PhyML 3.1 (Guindon et al., 2010).

## Supplementary Figures

**Figure S1.**
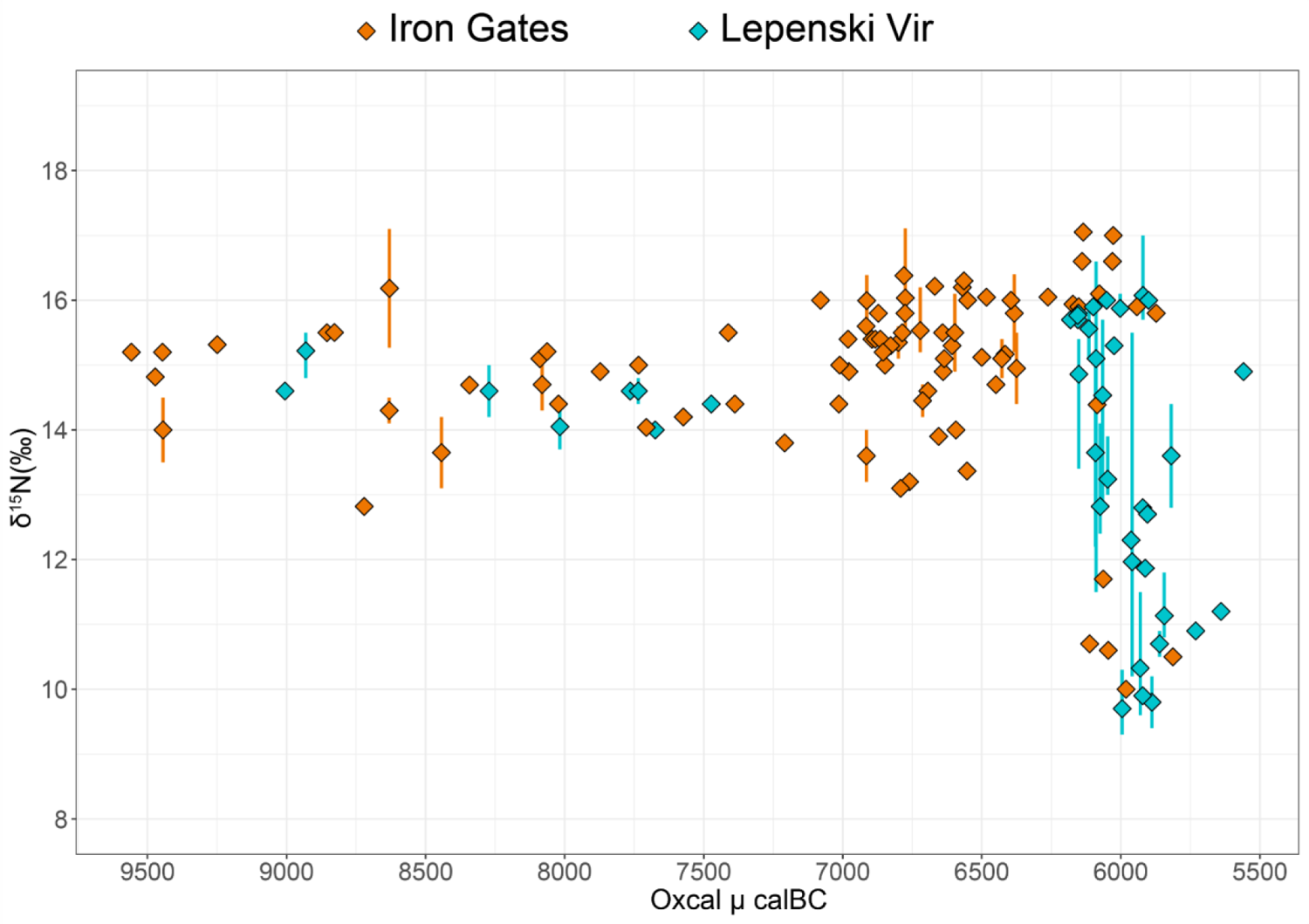
Shift in δ^15^N values observed in directly-dated humans from the Iron Gates (Oxcal μ). Any δ**^15^**N value above 10‰ is likely to indicate some intake of aquatic proteins. Based on previously reported ^14^C dates and δ**^15^**N values (full references in **Electronic Supplementary Material**). When multiple isotopic values are available from the same individual, both the median and the range are given. Excludes children <10.

**Figure S2.**
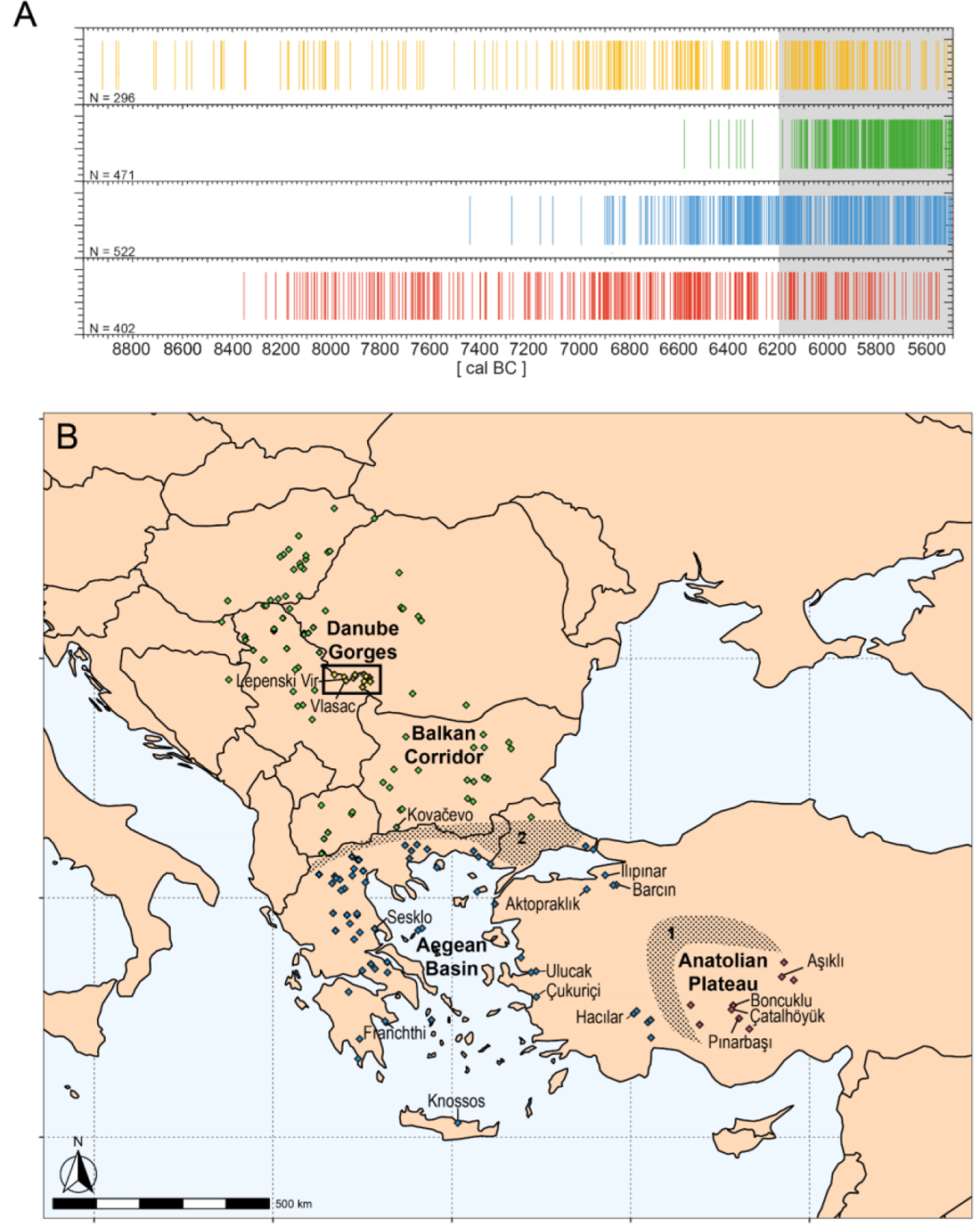
(A) Overview of ^14^C ages (total N=1,691) for 174 archaeological sites in four regions of Anatolia and Southeast Europe, with calibrated ages plotted by the Barcode Method (Weninger et al. 2014) in CalPal (Weninger & Jöris, 2008), using the IntCal09 calibration curve (Reimer et al., 2009). Each small vertical line shows the median value of the corresponding calibrated ^14^C-age. For the Anatolian Plateau (red), the Aegean Basin (blue) and the Balkan Corridor (green), only ^14^C dates associated with Neolithic layers (where food-production is evident or implied) are plotted. For the Danube Gorges (yellow), ^14^C dates associated with both Mesolithic and Neolithic occupations are plotted. Shading indicates the interval 6,200-5,500 BC - the approximate time frame of the main occupation at Lepenski Vir (Levels I-III); ^14^C dates as reported in the literature, including (Porčić et al., 2016), the CalPal Holocene and 14SEA databases (Reingruber & Thissen, 2016; Weninger, 2016); (B) distribution of the 174 radiocarbon-dated sites with ^14^C dates falling within the interval 9,500-5,500 BC. Geographic coordinates from the literature. The location of major agricultural frontier zones is indicated by 1 and 2 (tentative). Important sites mentioned in the text are listed here.

**Figure S3.**
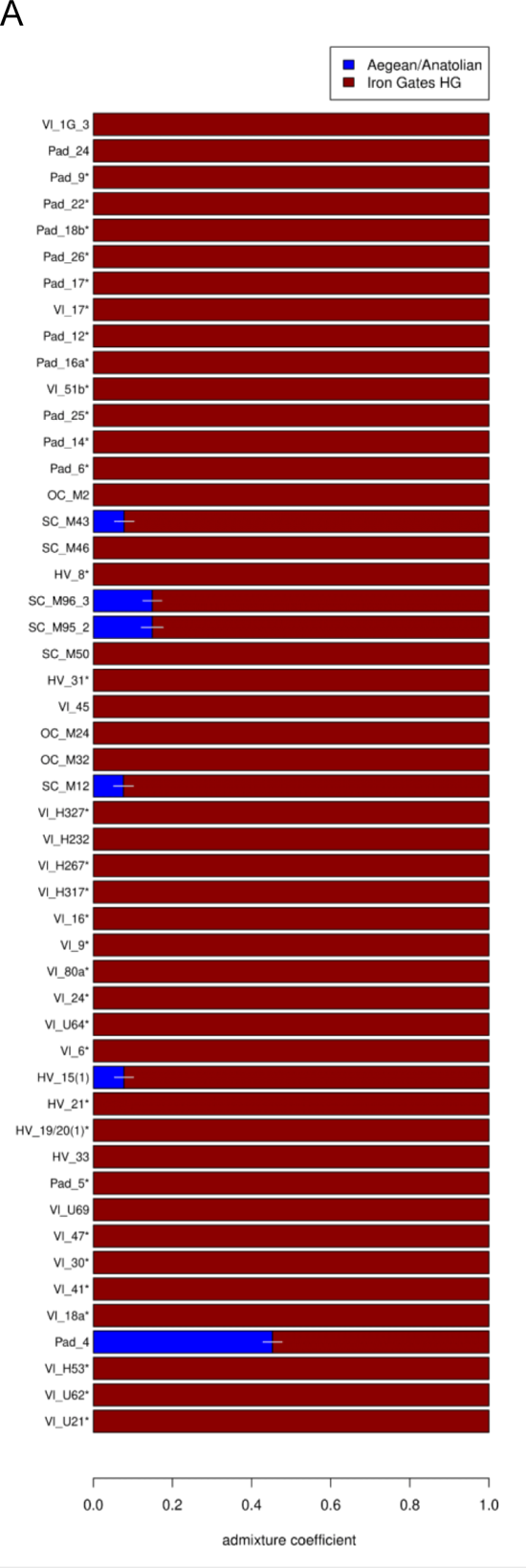
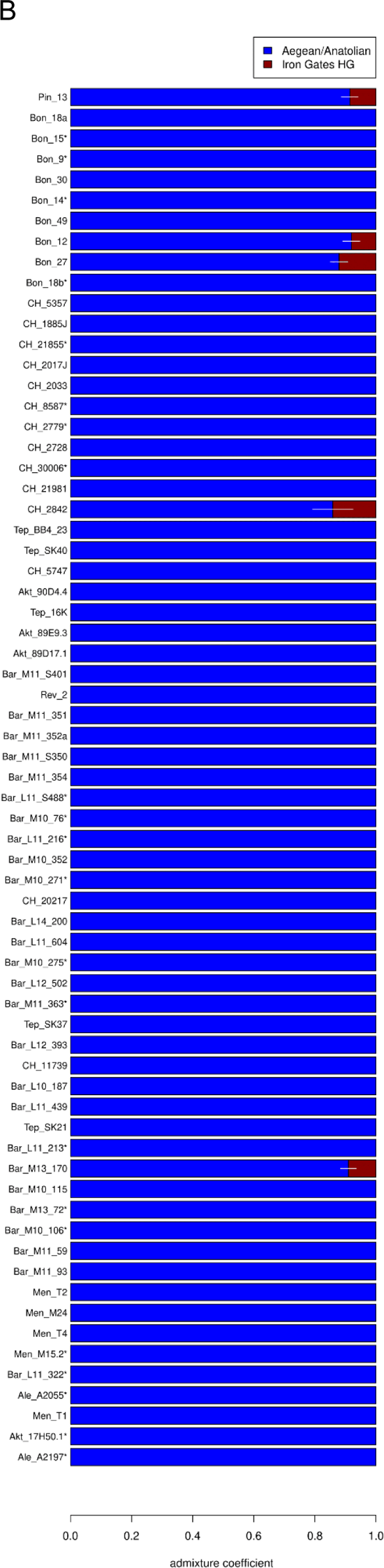
Iron Gates HG and Aegean-Anatolian ancestry components of Iron Gates (A; excluding Lepenski Vir) and Anatolian/Aegean populations (B) inferred with *qpAdm*. White error bars for admixed individuals show standard errors as provided by *qpAdm*.

**Figure S4.**
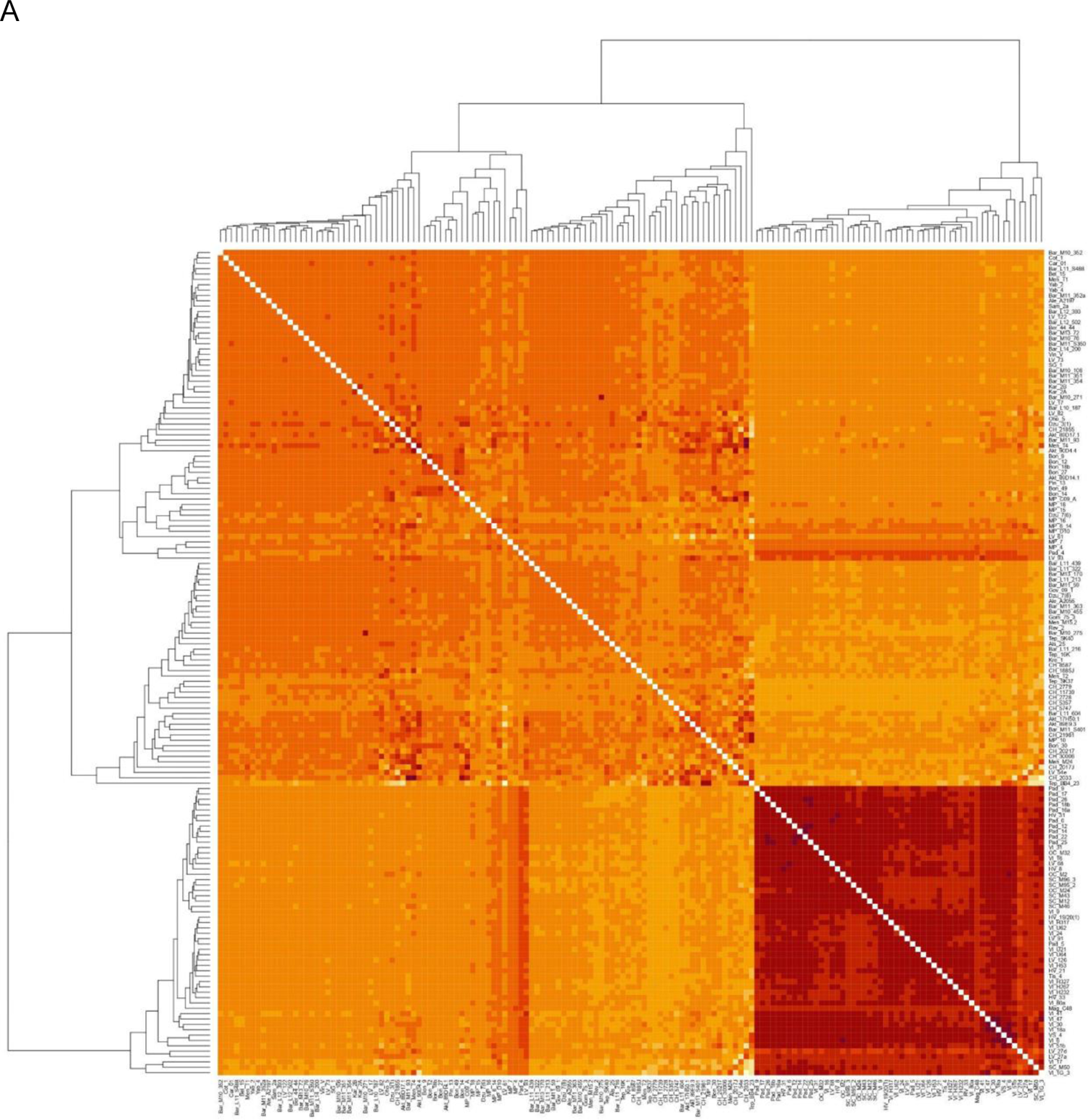
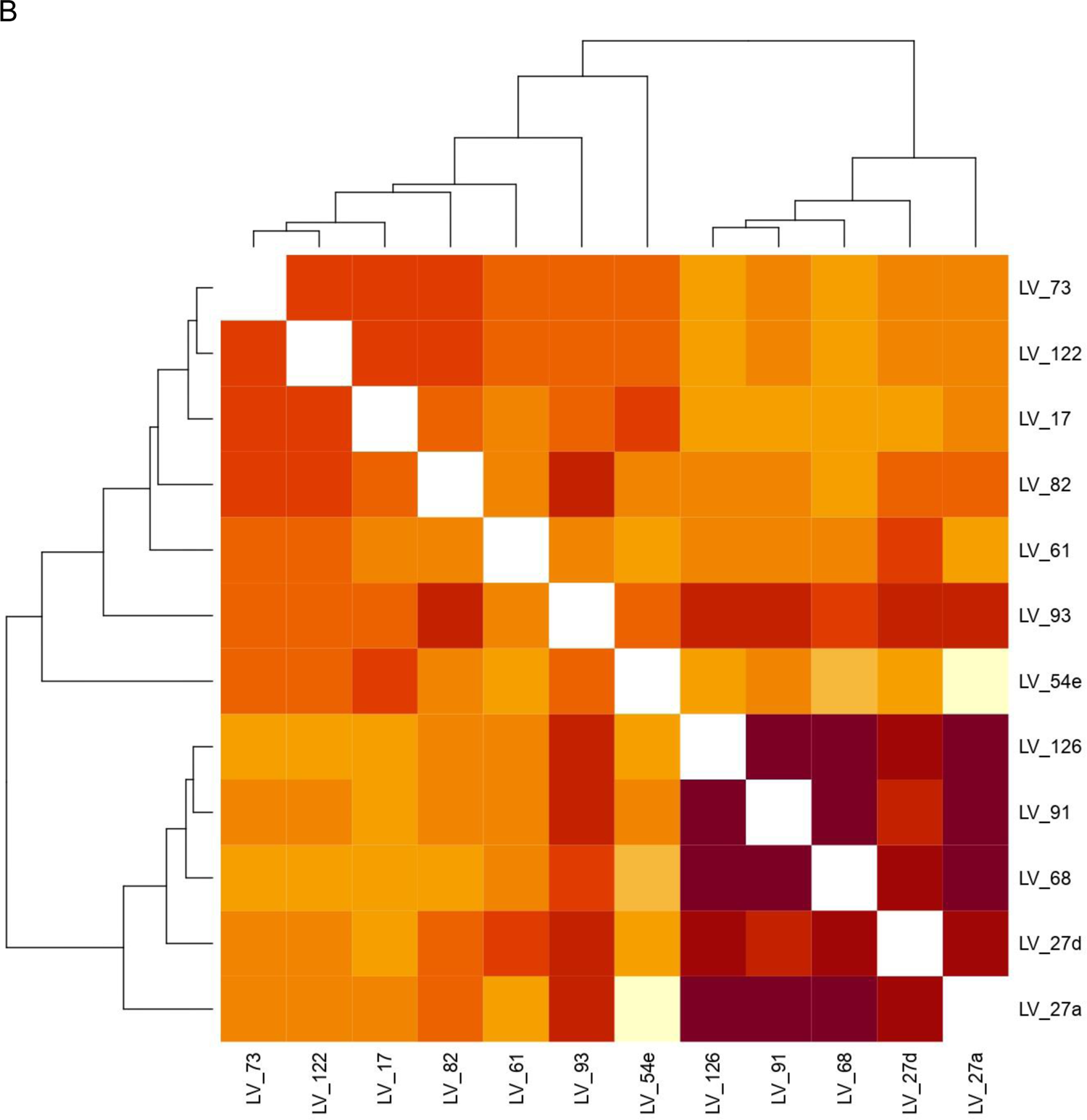
Pairwise outgroup *f_3_* similarities between Anatolian, Aegean and Iron Gates (A) and Lepenski Vir individuals (B).

**Figure S5.**
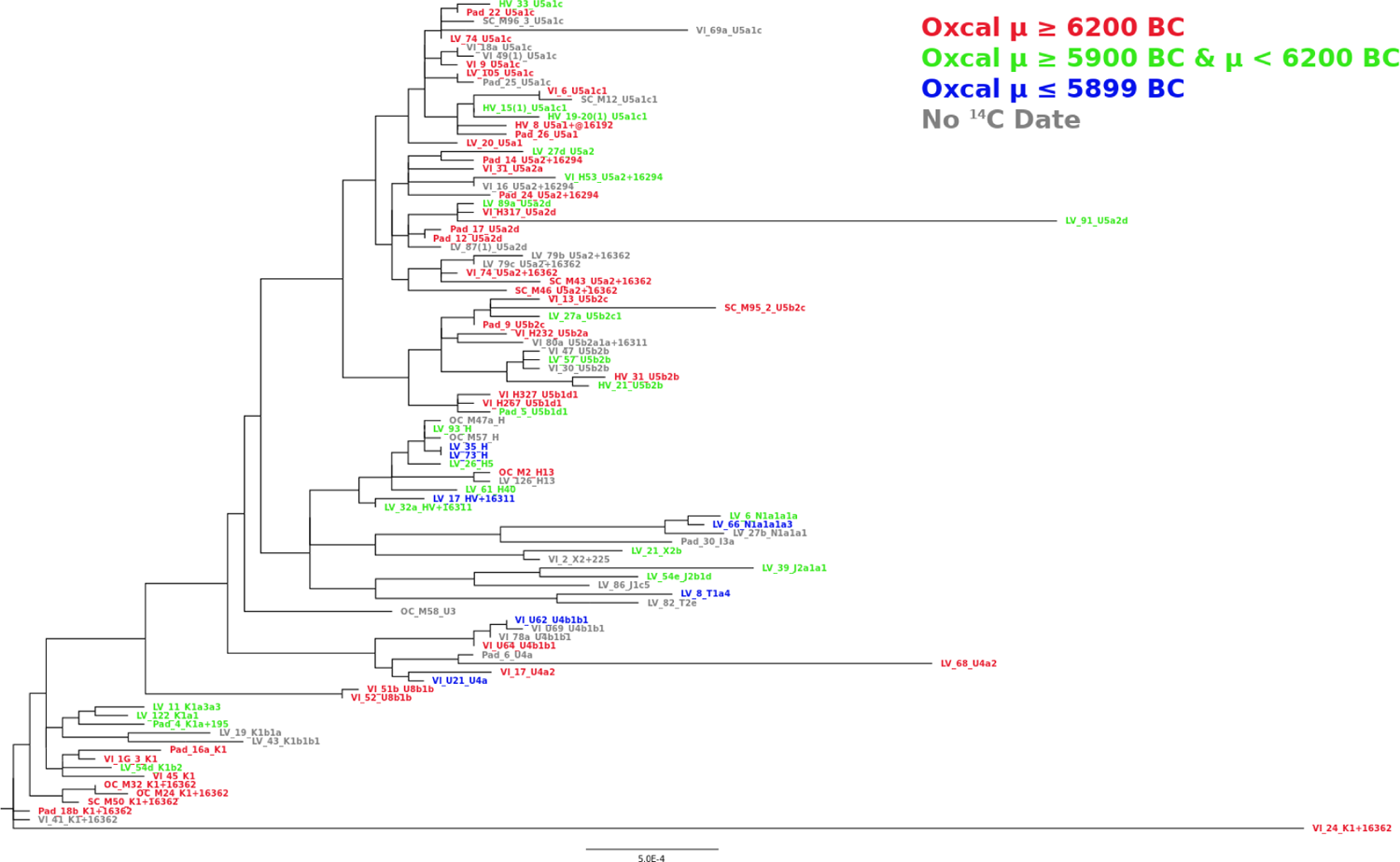
Tree built from MT-DNA from Mesolithic and Neolithic individuals of the Iron Gates. Tree was built with PhyML 3.1 (Guindon et al., 2010) on fasta sequences extracted from majorityAllele calls provided by ATLAS (Link et al., 2017).

**Figure S6.**
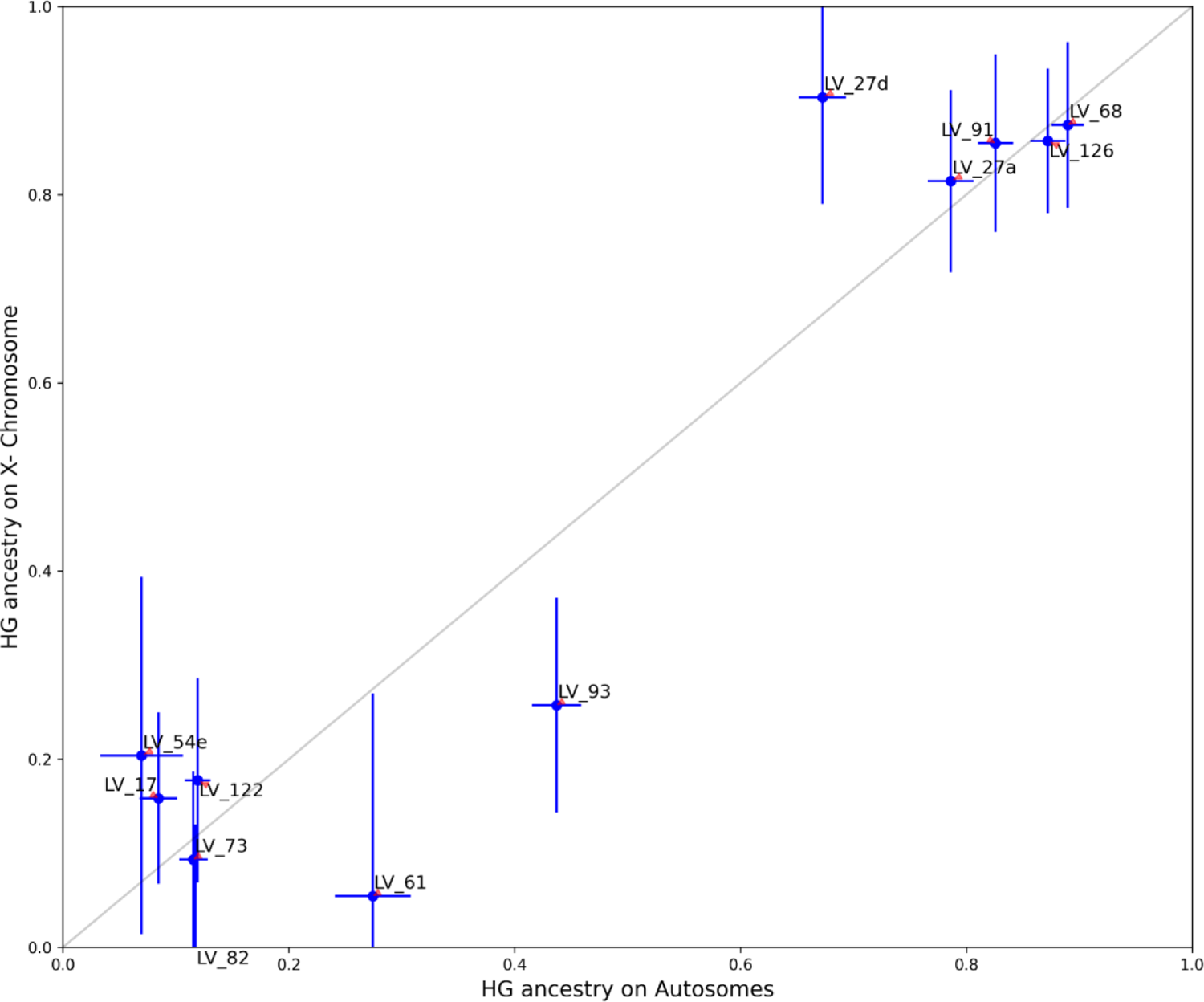
Iron Gates Hunter-Gatherer (HG) ancestry proportion on the X chromosome and the autosomes for Lepenski Vir individuals. Error bars represent 95% confidence intervals, see **Materials and Methods** for details.

**Figure S7.**
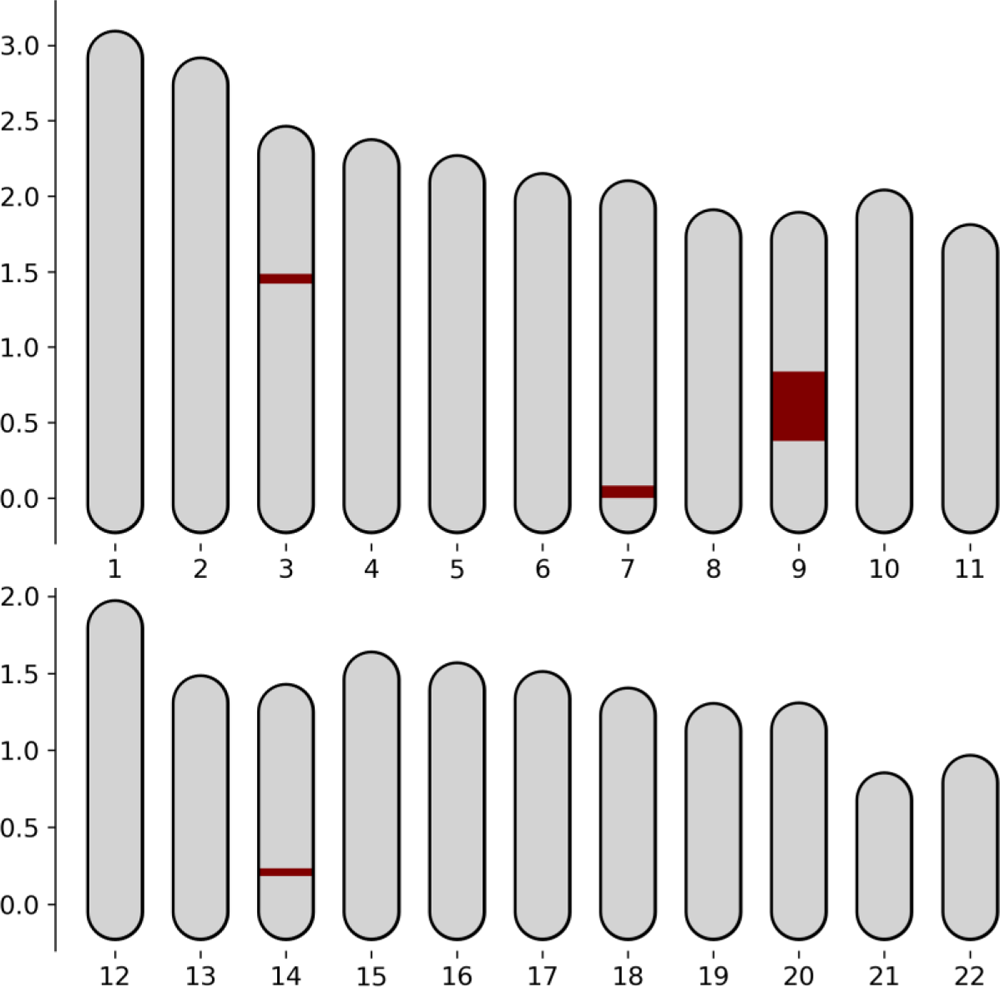
Runs of Homozygosity (>4cM) in Burial 73 (LEPE52) inferred by hapROH (Ringbauer et al., 2021).

**Figure S8.**
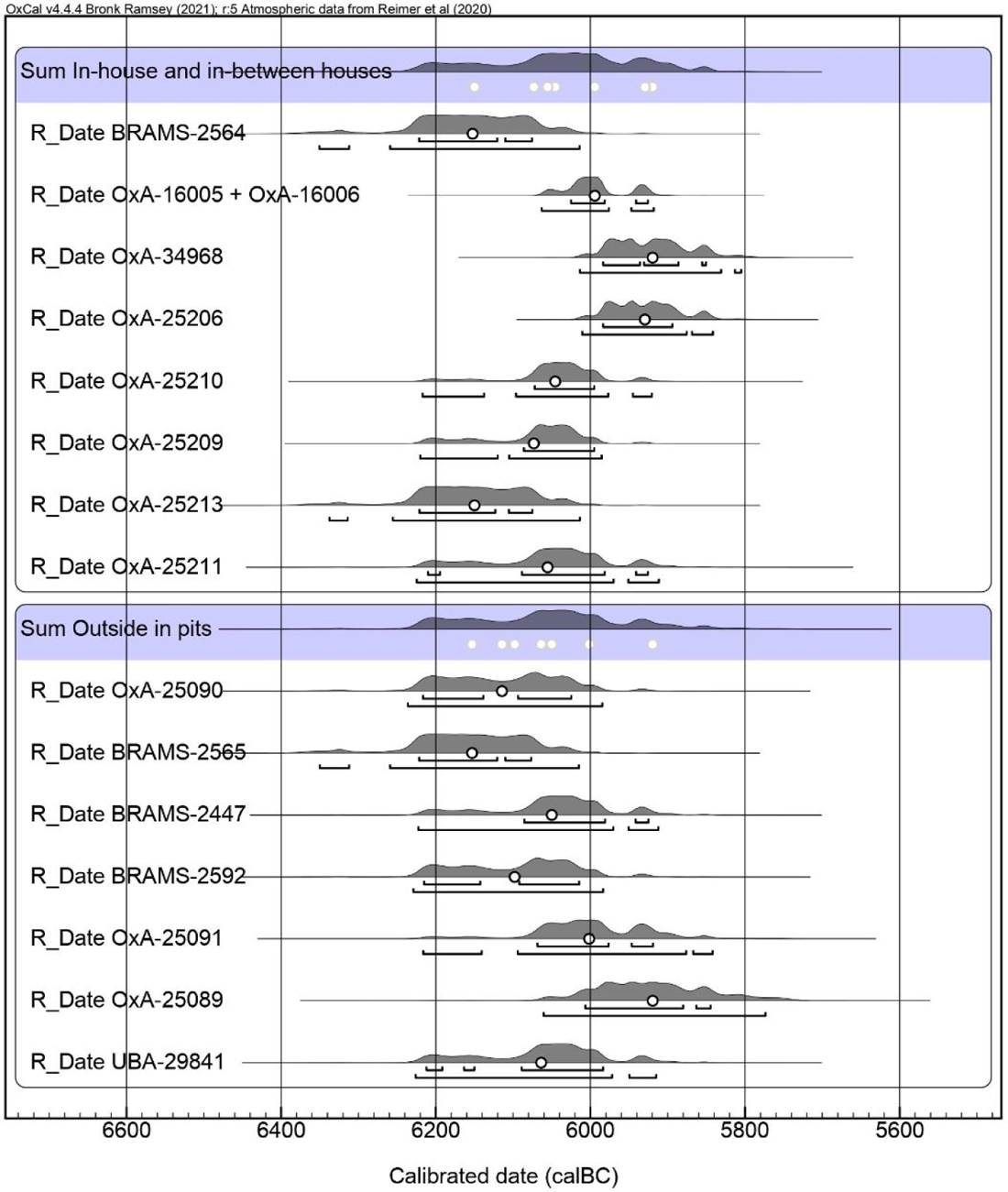
Summed probability distributions of calibrated ^14^C dates associated with burials in-house and in-between houses (top) and outside in pits (bottom). Supporting information and references in **Electronic Supplementary Material**.

**Figure S9.**
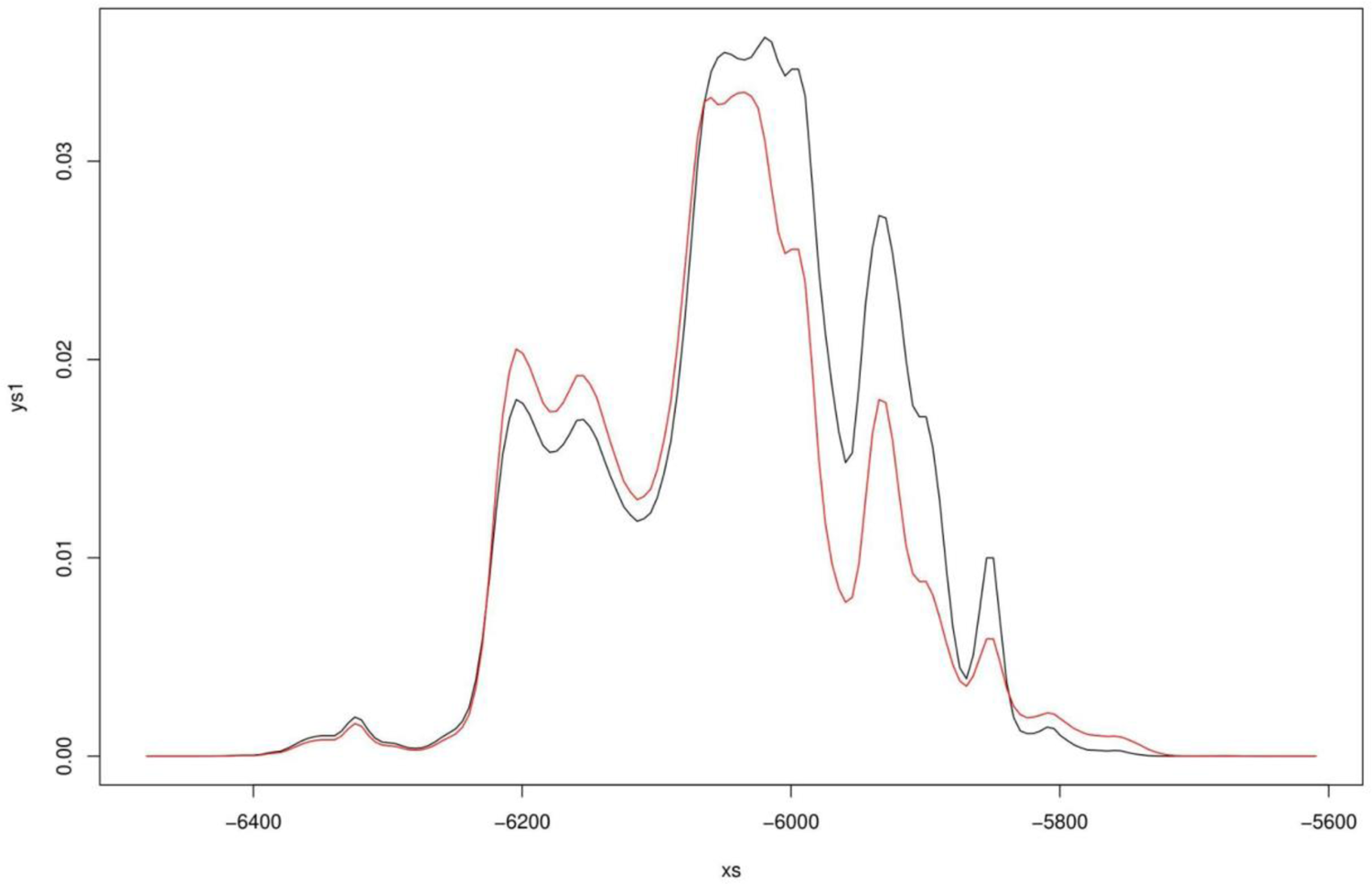
Summed probability distributions of calibrated ^14^C dates associated with burials in-house and in-between houses (black) and outside in pits (red). A two-sample Kolmogorov-Smirnov test does not reject the null hypothesis at significance level 5% that these are from the same distribution. See **Materials and Methods** for details on the KS-test used.

**Figure S10.**
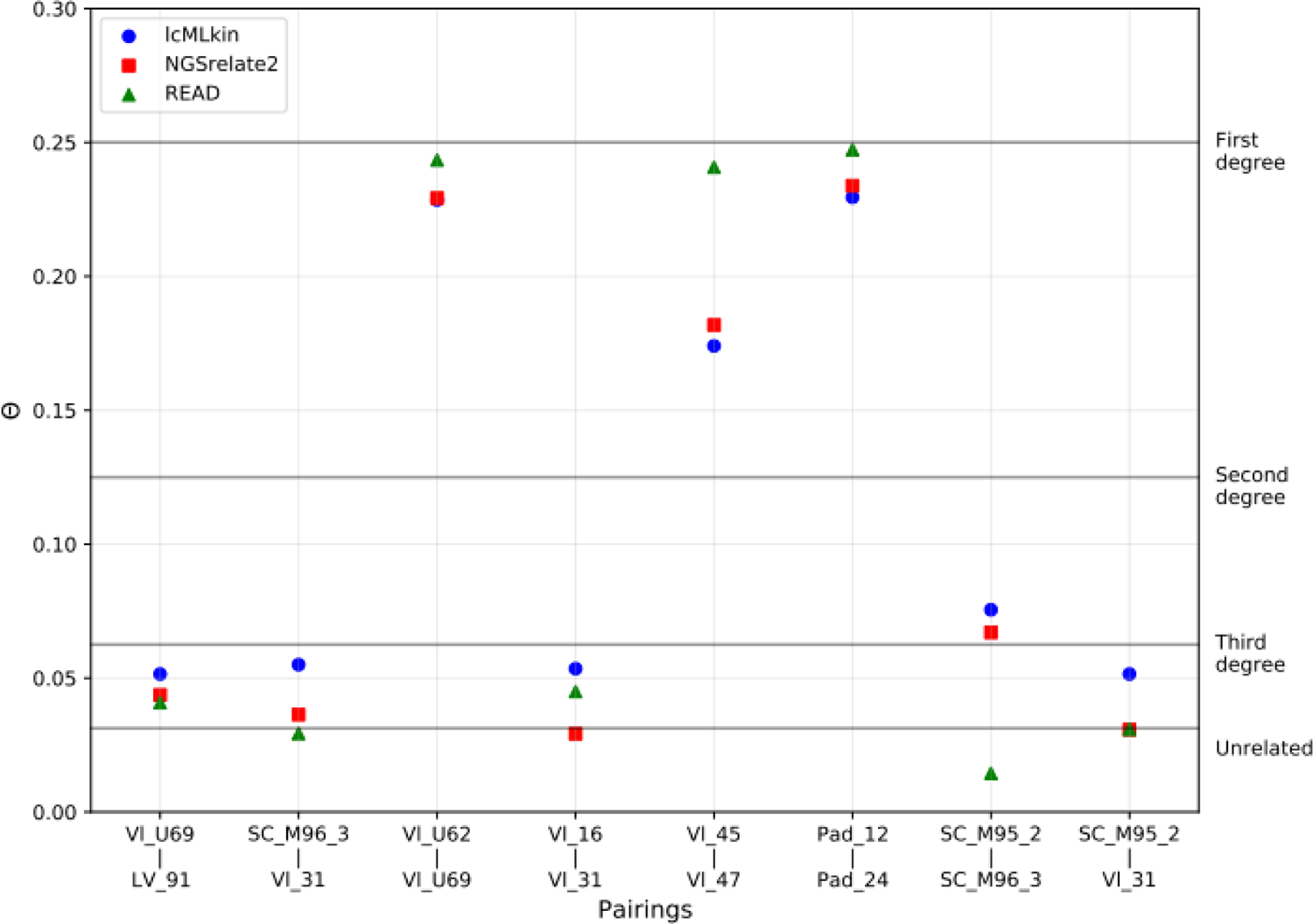
Kinship coefficient (θ) between pairs of individuals estimated by three different programs (lcMLkin, READ, ngsRelate2). Expected values for θ for first, second and third degree relatives are indicated by the black horizontal lines. Pairings were included if one method led to values close to or above third degree relation.

**Figure S11.**
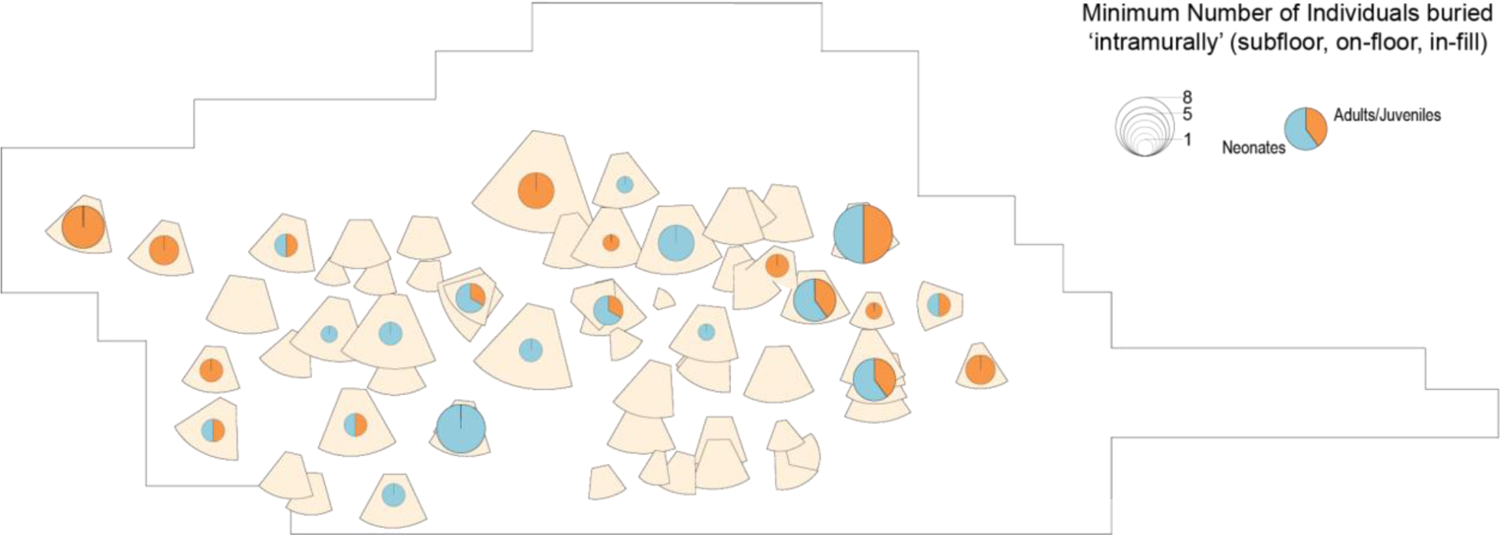
Distribution of ‘intramural’ burials at Lepenski Vir for all building phases (Levels I-III). ‘Burials’ refers to discrete individuals, regardless of completeness and burial practice (for a full description of burials at Lepenski Vir see (Borić, 2016)). Not all burials relate to the occupation of the trapezoidal houses, which were abandoned in Level III. As some houses were reconstructed multiple times in the same location, the total number of burials in relation to each house sequence is indicated.

**Figure S12.**
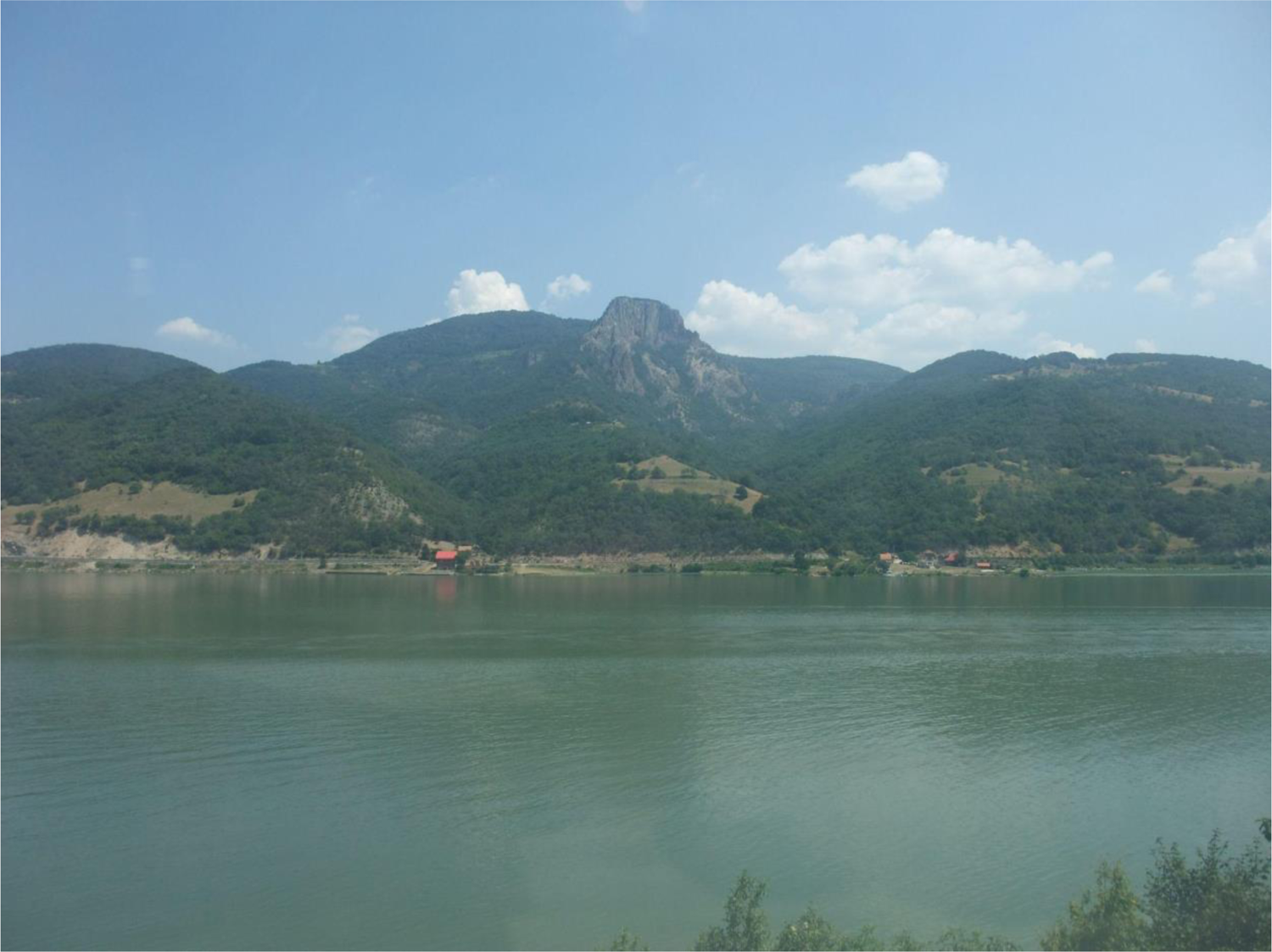
The distinctive Treskavac Mountain on the opposite side of the river (photograph: Maxime Brami)

